# Metallothionein 3-zinc axis suppresses caspase-11 inflammasome activation and impairs antibacterial immunity

**DOI:** 10.1101/2021.07.28.454033

**Authors:** Debabrata Chowdhury, Jason C Gardner, Abhijit Satpati, Suba Nookala, Santhosh Mukundan, Aleksey Porollo, Julio A. Landero Figueroa, Kavitha Subramanian Vignesh

## Abstract

Non-canonical inflammasome activation by mouse caspase-11 (or human CASPASE- 4/5) is crucial for the clearance of certain gram-negative bacterial infections, but can lead to severe inflammatory damage. Factors that promote non-canonical inflammasome activation are well recognized, but less is known about the mechanisms underlying its negative regulation. Herein, we identify that the caspase-11 inflammasome in mouse and human macrophages (Mϕ) is negatively controlled by the zinc (Zn^2+^) regulating protein, metallothionein 3 (MT3). Upon challenge with intracellular lipopolysaccharide (iLPS), Mϕ increased MT3 expression that curtailed the activation of caspase-11 and its downstream targets caspase-1 and interleukin (IL)-1β. Mechanistically, MT3 increased intramacrophage Zn^2+^ to downmodulate the TRIF-IRF3-STAT1 axis that is prerequisite for caspase-11 effector function. MT3 suppressed activation of the caspase-11 inflammasome, while caspase-11 and MT3 synergized in impairing antibacterial immunity. The present study identifies an important yin-yang relationship between the non-canonical inflammasome and MT3 in controlling inflammation and immunity to gram-negative bacteria.

**GRAPHICAL ABSTRACT:** 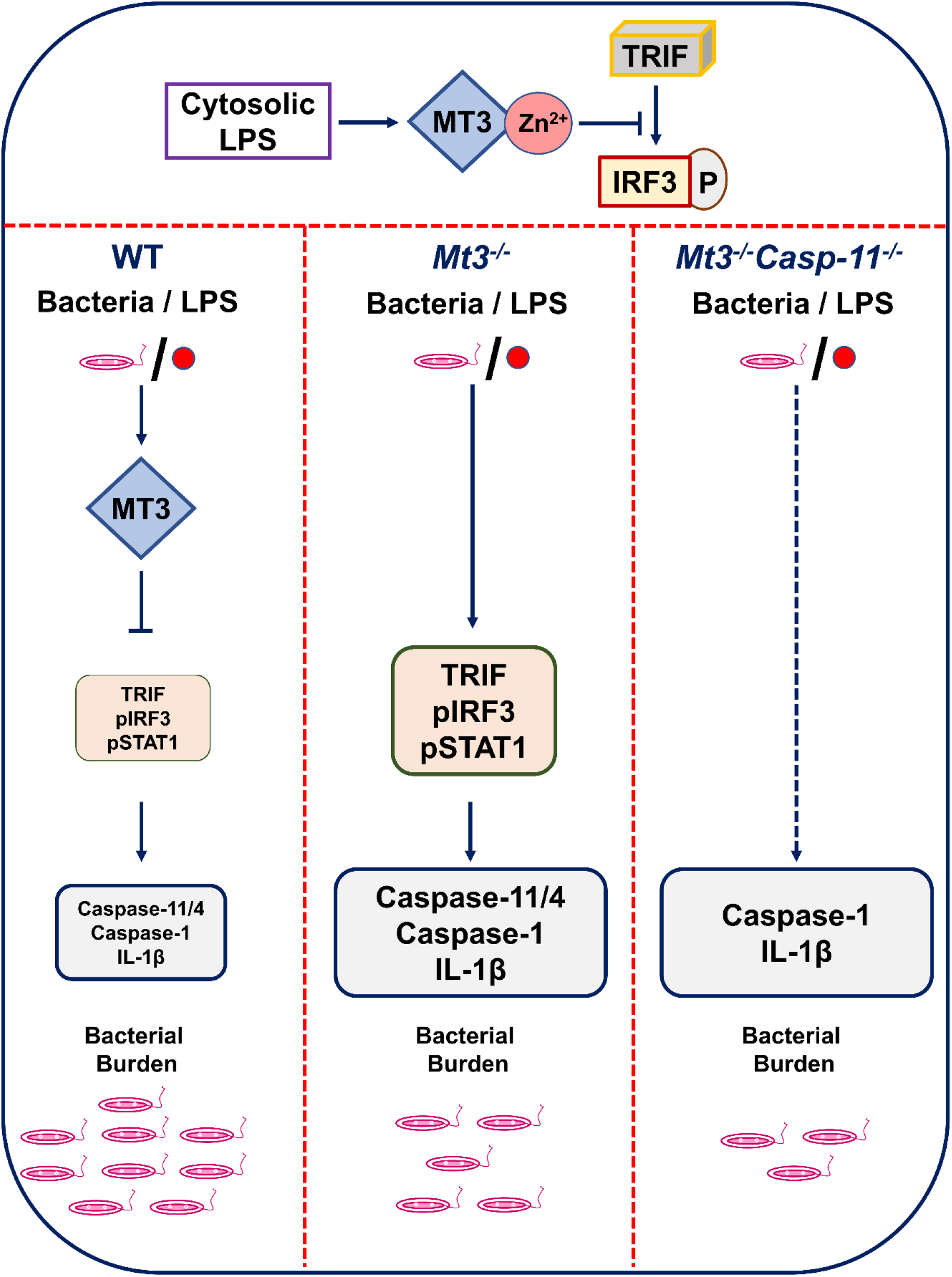

## Introduction

Gram-negative bacteria that cause more than 30% of the total healthcare-associated infections worldwide remain a global concern of morbidity and mortality ^1^. Assembly of inflammasome complexes during bacterial pathogenesis drives robust inflammation and shapes antibacterial immune responses. The non-canonical inflammasome is activated when innate immune cells such as Mϕ sense bacterial ligands in the cytosol ^2^. LPS from bacterial cell walls enters the cytosol during bacterial escape from vacuoles or via rupture of outer membrane vesicles (OMV) ^3^. iLPS directly binds caspase-11, triggering the non-canonical inflammasome cascade, followed by activation of pro-caspase-1 to caspase-1, processing of pro-IL-1β to mature IL-1β and pyroptosis, a lytic form of programmed cell death. Activation of gasdermin D (GSDMD) by caspase-11 and caspase-1, leads to pore formation on the cell membrane facilitating the exit of IL-1β from Mϕ ^4^. Unrestricted activation of this inflammatory cascade is a major underlying cause of tissue damage and sepsis-associated mortality ^2^. Thus, it is crucial to thoroughly understand the molecular cues that guard against excessive activation of the caspase-11 inflammasome. Although factors that promote non-canonical inflammasome activation have been extensively studied, less is known about the mechanisms that negatively regulate it.

MTs are Zn^2+^ regulating proteins induced by endogenous and exogenous stimuli including cytokines, infection, oxidative stress and heavy metals ^5, 6^. Intracellular availability of total Zn^2+^, exchangeable Zn^2+^ and Zn^2+^ redistribution among proteins is tightly regulated by MTs ^7^. Mice have 4 MT isoforms (MT1-4), whereas >16 MT isoforms are present in humans ^8^. Our knowledge on the role of the MT family in immune responses largely emerges from studies on MT1 and MT2. We and others have shown that MT1 and MT2 promote antifungal and antibacterial immunity in Mϕ primarily via Zn^2+^ sequestration ^9, 10^. MT3, on the other hand, suppresses manifestation of proinflammatory phenotypic and metabolic changes and impairs antifungal immunity *in vitro* in Mϕ and *in vivo*. Mϕ expression of MT3 is inducible by IL-4 stimulation. In these cells, MT3 increases intracellular free-Zn^2+^, promotes Zn^2+^ uptake by a prototypic intracellular pathogen and favors microbial survival ^11^. Thus, MTs and their ability to regulate Zn^2+^ homeostasis intricately ties Mϕ inflammatory responses to antimicrobial defense.

Zn^2+^ is indispensable in many biochemical processes due to its role in structural and catalytic functions of enzymes and macromolecules. Zn^2+^ excess or deficiency compromises the development and function of immune cells including monocytes and Mϕ, leading to increased risk of infection ^12^. Zn^2+^ also has a profound role as a signaling ion attributable to the transient changes in intracellular exchangeable Zn^2+^. LPS triggers increased Zn^2+^ import in leukocytes, monocytes and Mϕ ^13, 14^. In peripheral blood mononuclear cells (PBMCs) and human Mϕ (hMϕ), LPS-induced Zn^2+^ influx promotes IL- 1β production ^14^. In contrast, exogenous exposure to Zn^2+^ in human monocytes may reduce IL-1β production due to inhibition of nucleotide phosphodiesterases ^15^. Thus, the effects of Zn^2+^ on signaling and cytokine production are context and Zn^2+^ concentration- dependent. Changes in ion flux, specifically K^+^ and Cl^-^ egress and intracellular mobilization of Ca^2+^ underlie canonical nod-like receptor pyrin domain containing-3 (NLRP3) inflammasome activation ^16^. Intriguingly, Zn^2+^ exerts disparate effects on activation of this cascade. Long-term Zn^2+^ depletion disrupts lysosomal integrity leading to increased activation of the canonical NLRP3 inflammasome ^17^. On the other hand, short-term chelation of Zn^2+^ attenuates the canonical pathway due to impaired function of the pannexin-1 receptor ^18^.

The importance of Zn^2+^ regulation by MTs in the non-canonical inflammasome pathway remain unexplored. Given the suppressive role of MT3 in Mϕ inflammatory responses ^19^, we hypothesized that MT3 negatively regulates the highly inflammatory caspase-11 activation cascade. Bioinformatics analysis predicted the involvement of MT3 in regulating non-canonical inflammasome-associated pathways. Using a combination of protein-protein interaction network analysis, immunological and mass-spectrometric approaches, we demonstrate that triggering caspase-11 activation results in a profound, gradual increase in the Mϕ Zn^2+^ pool mediated by MT3. The increase in Zn^2+^ attenuates signaling via toll/interleukin-1 receptor (TIR) domain containing adaptor-inducing interferon (IFN)β - interferon regulatory factor 3 - signal transducer and activator of transcription factor 1 (TRIF-IRF3-STAT1), a pathway that is prerequisite for caspase-11 inflammasome activation ^20^. Zn^2+^ deficiency augments, whereas Zn^2+^ supplementation suppresses the non-canonical inflammasome in Mϕ. Using whole-body *Mt3^-/-^* and myeloid-MT3-deficient mice, we elucidate that MT3 blunts non-canonical inflammasome activation *in vitro* and *in vivo* upon challenge with iLPS or gram-negative bacteria but not gram-positive bacteria. Importantly, this function of MT3 is conserved in hMϕ. Although caspase-11 and MT3 form a negative regulatory loop, we find that these two molecules synergize in compromising antibacterial immunity. Our data uncover a previously unknown yin-yang relationship whereby the MT3-Zn^2+^ axis exerts a brake on non-canonical inflammasome activation but the functions of MT3 and caspase-11 converge in crippling immunity to invading bacteria.

## Results

### MT3 suppresses activation of the non-canonical inflammasome *in vitro*

MT3 attenuates cell death in neuronal and glial cells, but the precise underlying mechanisms are not fully understood ^21^. As non-canonical inflammasome activation leads to pyroptotic cell death, we investigated if MT3 effector function is related to caspase-11 activation in Mϕ. We explored whether MT3 is involved in inflammatory and cell-death processes using functional enrichment analysis. We assessed protein-protein interaction networks of MT3 in *Mus musculus* and *Homo sapiens* using the STRING database ^22^. The MT3 interaction partners significantly enriched 15 mouse and 43 human gene ontology categories for biological processes (GO BP) related to programmed cell death (PCD), LPS responses, signaling via TRIF, IL-1 and type-I IFN, cytokine responses and several immune processes. A complete network of MT3 interactions and GO BP categories is in **Table 1 and Extended Data Table 1, Supplementary Files 1 and 2**. PCD and LPS responses are linked to inflammasome activation and more specifically, TRIF and type-I IFN signaling are tied to the non-canonical inflammasome pathway. A lack of TRIF signaling ablates non-canonical inflammasome activation in response to iLPS without impacting Mϕ response to canonical NLRP3 triggers such as ATP and nigericin ^20^. Thus, our bioinformatics analysis together with our previously reported role for MT3 in suppressing proinflammatory responses in Mϕ led us to investigate whether MT3 negatively regulates the non-canonical inflammasome pathway.

**Table 1.**
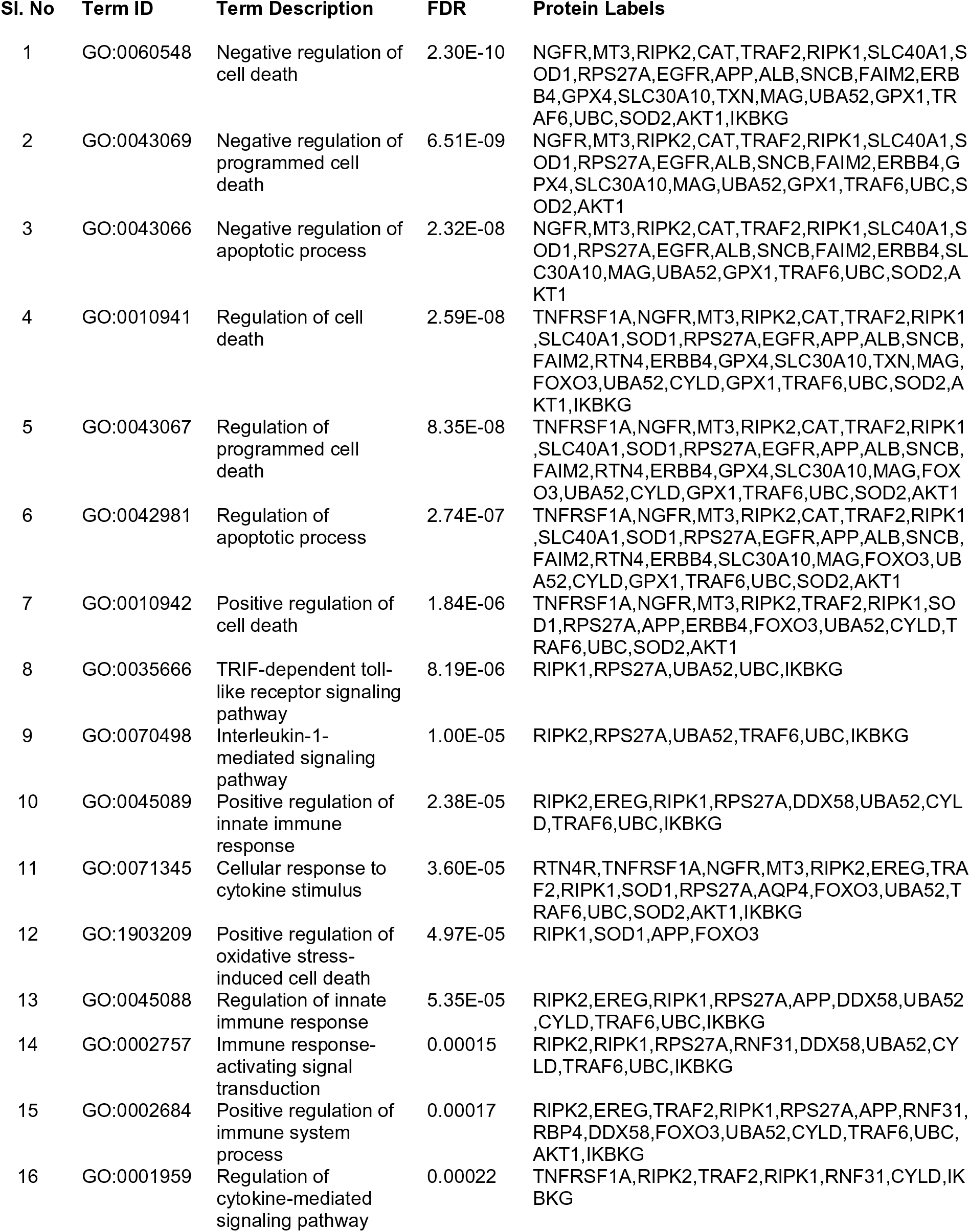
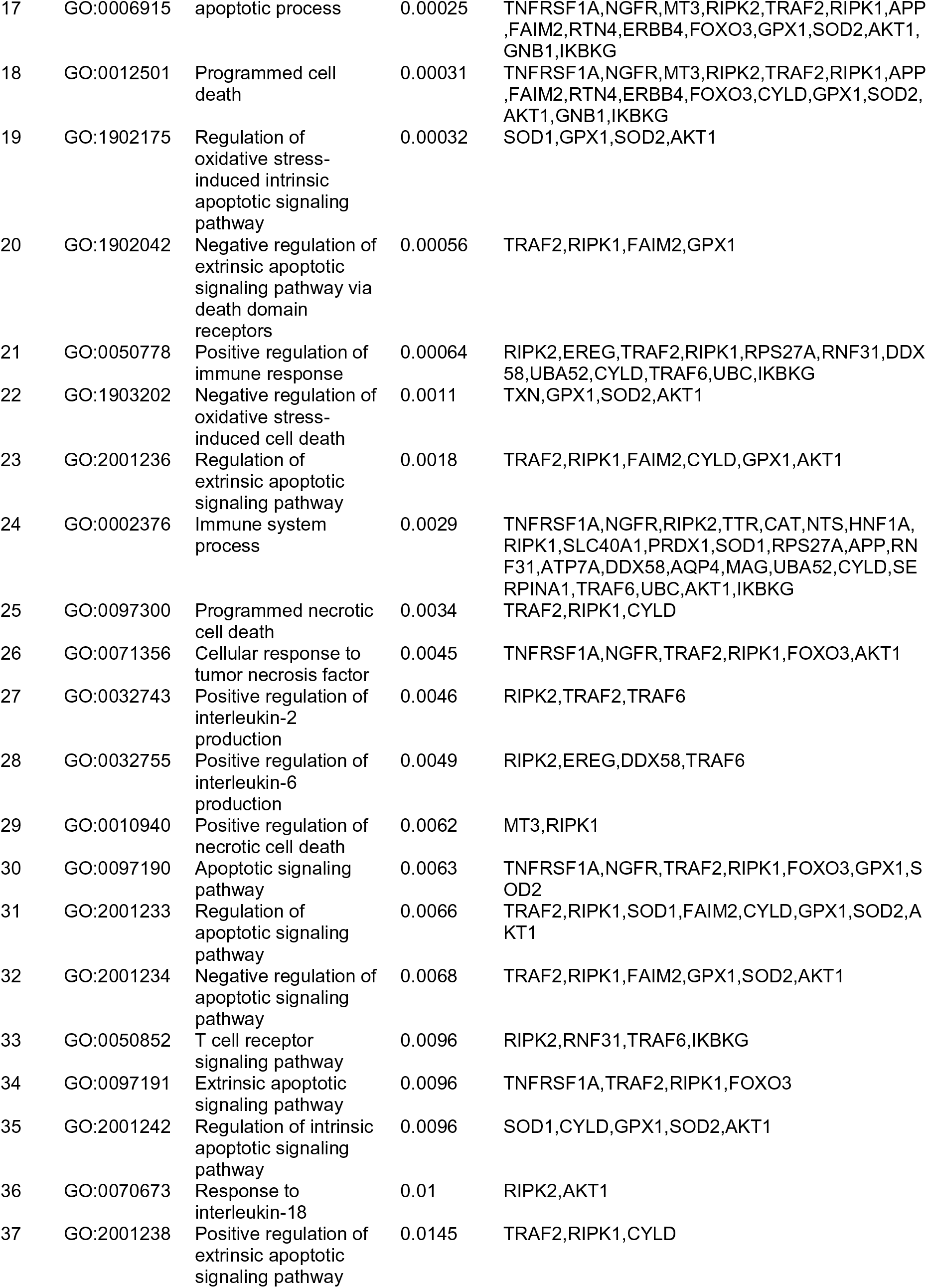
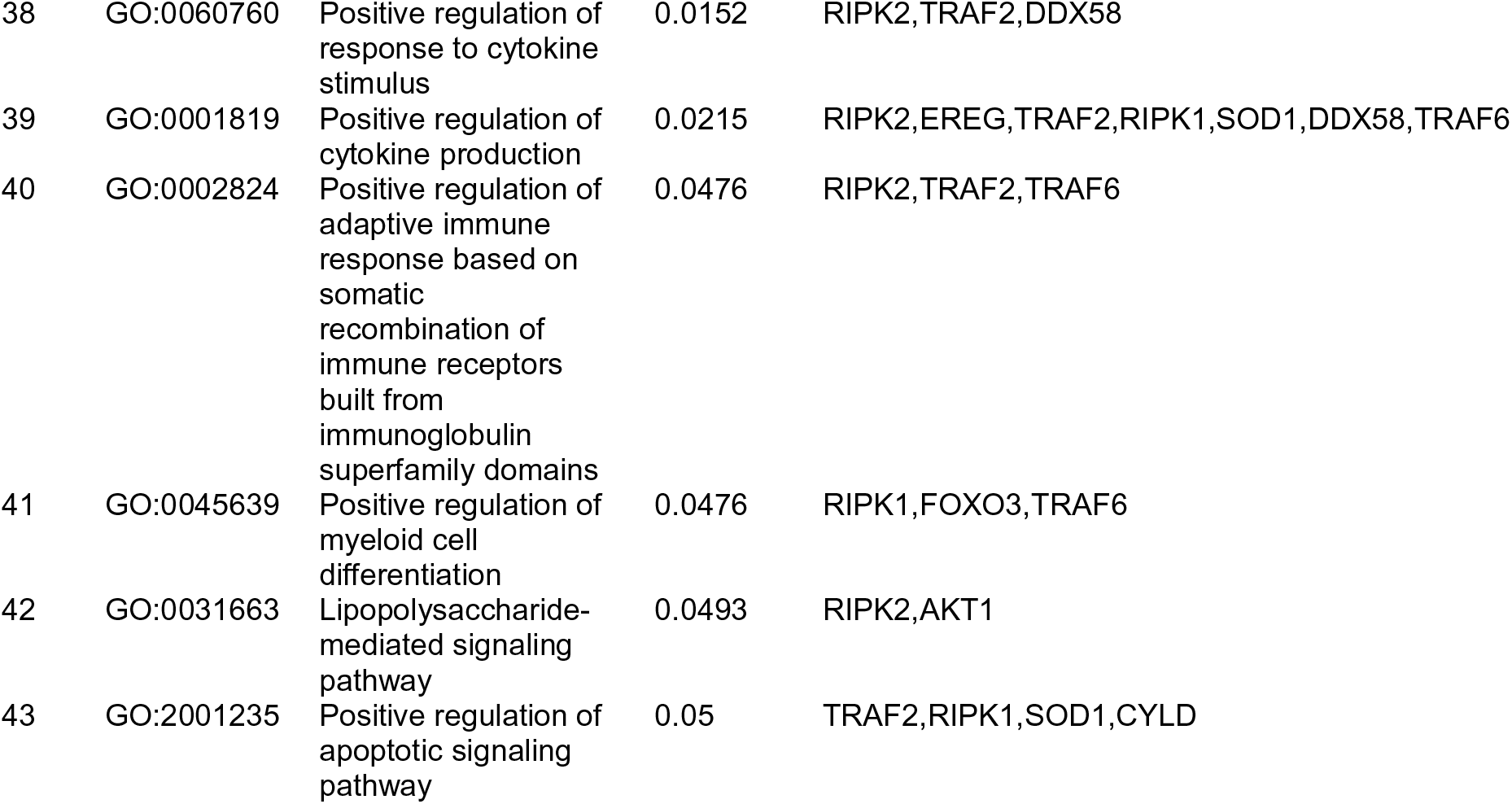
Protein interaction network of *Mus musculus* MT3 to determine functionally enriched GO BP categories using the STRING database. **See also Extended Data Table 1 and Extended Data Files 1,2 |**

In Mϕ, exposure to iLPS triggers caspase-11 activation and cell death by pyroptosis ^2^. We directly assessed if MT3 regulates non-canonical inflammasome activation using WT and *Mt3^-/-^* mice. BMDMϕ were exposed to iLPS or vehicle control and time-dependent changes in the gene expression of *Mt1*, *Mt2* and *Mt3* were examined. *Mt3* expression increased gradually from 1 hour (h) and peaked at 48h in WT BMDMϕ challenged with iLPS **(Fig. 1A)**. The expression of *Mt1* and *Mt2* peaked at 6h, but receded over time in both WT and *Mt3^-/-^* BMDMϕ **(Extended Data Fig. 1A and 1B)**. We examined if MT3 regulated non-canonical inflammasome activation by assessing pro- and active forms of caspase-11, caspase-1 and IL-1β in WT and *Mt3^-/-^* BMDMϕ challenged with iLPS. A lack of MT3 exacerbated activation of caspase-11, caspase-1 and IL-1β. Pro-caspase- 11, pro-caspase-1 and pro-IL-1β proteins were similar between iLPS treated WT and *Mt3^-/-^* BMDMϕ **(Fig. 1B)**.

**Fig. 1.**
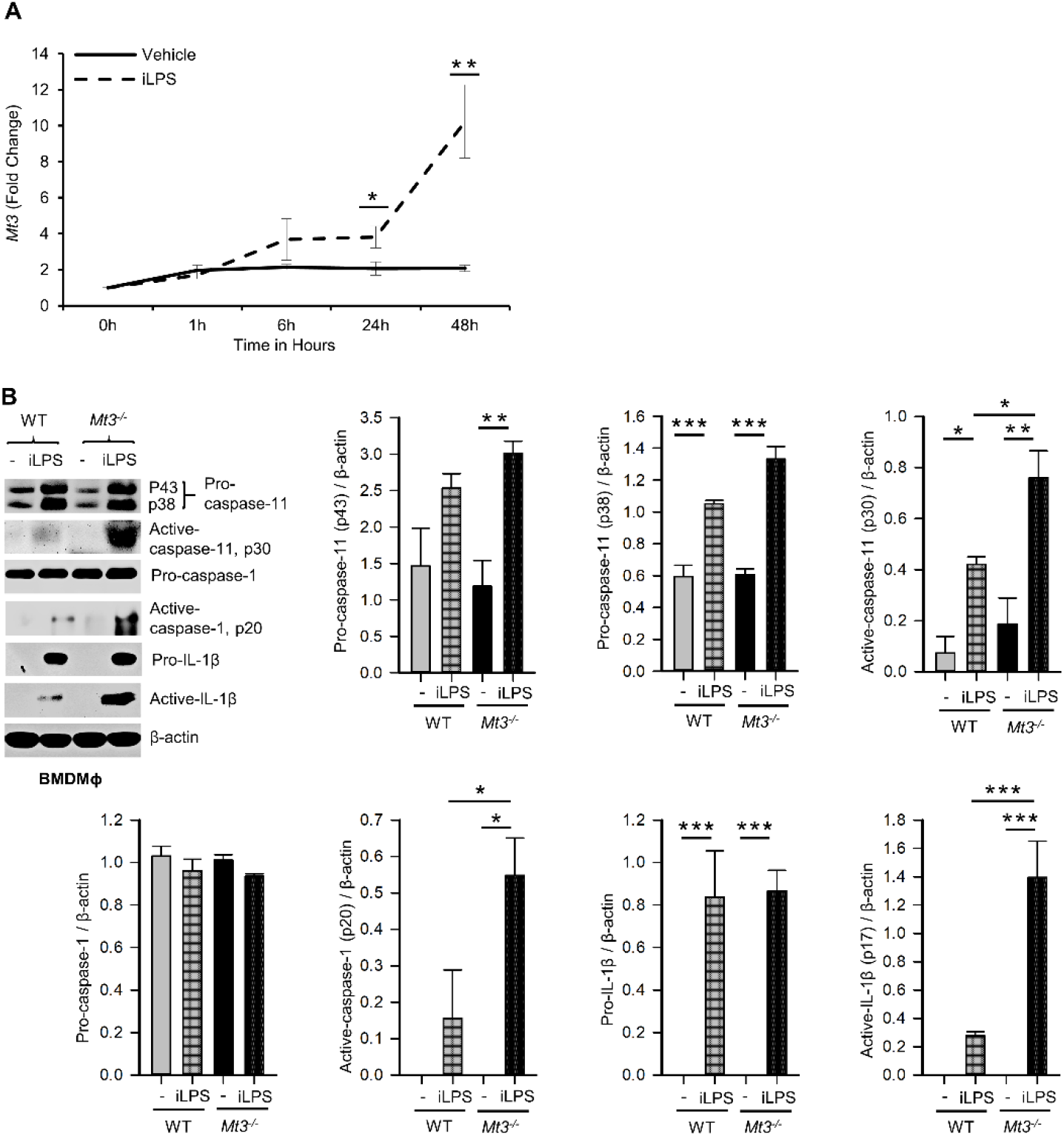
MT3 suppresses caspase-11 inflammasome activation in BMDMϕ. **See also Extended Data Fig. 1 | A**, qRT-PCR analysis of *Mt3* expression in WT BMDMϕ stimulated with iLPS (2 μg/ml) or vehicle control, 3-5 independent experiments, two-tailed t-test. **B**, Western Blots of pro- and active-caspase-11, pro-caspase-1, pro-IL1β and β-actin in cell lysates and active-caspase-1 and active-IL-1β in supernatants of WT and *Mt3^-/-^* BMDMϕ stimulated with iLPS (10 μg/ml) or vehicle for 48h. Bar graphs are densitometric analysis of targets normalized to β-actin, 3-4 independent experiments, one-way ANOVA, data are mean ± SEM.

### MT3 represses CASPASE-4 activation and antibacterial resistance in human Mϕ

Gram-negative bacteria activate the non-canonical inflammasome via CASPASE-4 in hMϕ ^23^. We investigated whether human MT3, similar to mouse MT3, suppressed non-canonical inflammasome activation and antibacterial immunity. Human monocyte-derived Mϕ obtained from PBMCs were transfected with scramble siRNA or *MT3* siRNA followed by transfection with iLPS **(Fig. 2A)**. MT3 deficiency resulted in elevated activation of CASPASE-4 and heightened release of IL-1β from hMϕ **(Figs. 2B and 2C)**. We previously showed that a lack of MT3 increased resistance of mouse BMDMϕ to *Escherichia coli* ^19^. We therefore queried whether silencing MT3 in hMϕ impaired bacterial clearance *in vitro*. hMϕ treated with scramble siRNA or *MT3* siRNA were infected with *E. coli* K12 for 24h. MT3-deficient hMϕ exerted a sharp decline in intracellular bacterial survival compared to control hMϕ **(Fig. 2D)**.

**Fig. 2.**
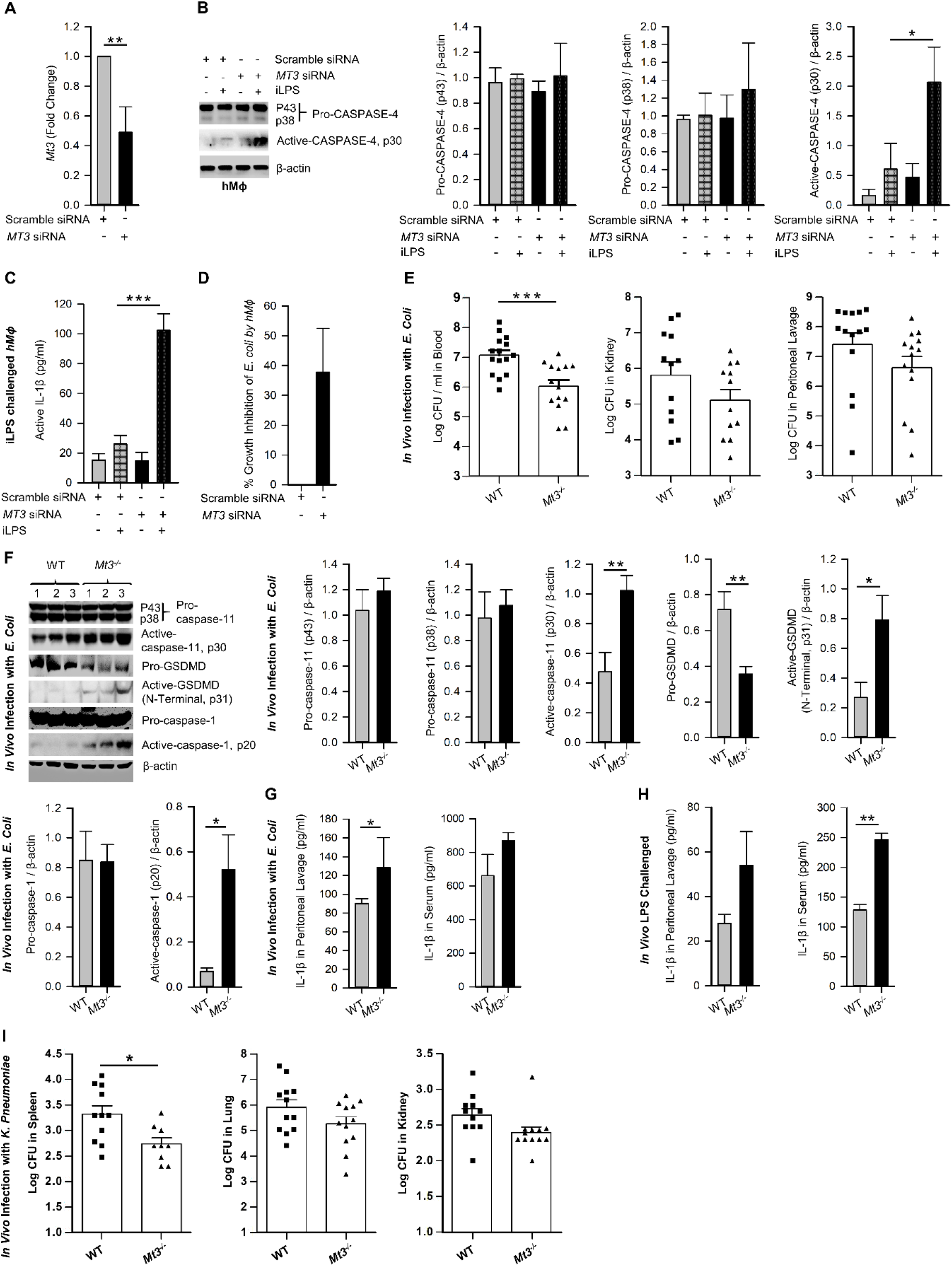
MT3 curtails CASPASE-4 and caspase-11 signaling and antibacterial immunity in hMϕ and *in vivo*. **See also Extended Data Fig. 2 | A**, *MT3* expression analyzed by qRT-PCR in hMϕ transfected with scramble siRNA or *MT3* siRNA for 24h, 3 independent experiments, two-tailed t-test. **B**, Scramble siRNA or *MT3* siRNA treated hMϕ stimulated with iLPS (10 μg/ml) or vehicle for 48h. Immunoblots of pro-CASPASE-4 and active-CASPASE-4 in cell extracts, 3 independent experiments, one-way ANOVA. **C**, Active-IL-1β measured by ELISA in supernatants of hMϕ treated as above, 3 independent experiments, one-way ANOVA. **D**, *E. coli* growth inhibition in hMϕ transfected with *MT3* siRNA and infected with 25 *E. coli* (K12): 1 hMϕ for 24h compared to scramble siRNA treated hMϕ, 3 independent experiments, two-tailed t-test. **E**, WT and *Mt3^-/-^* mice infected *i.p.* with 1 X10^9^ *E. coli* for 6h, log CFUs of *E. coli* in blood, kidney and peritoneal lavage samples, n = 12-15 per group, two-tailed t-test. **F**, Western blots of inflammasome mediators in kidney homogenates of WT and *Mt3^-/-^* mice infected as above, n = 6 per group, two-tailed t-test. **G**, WT and *Mt3^-/-^* mice infected *i.p.* with 1 X10^9^ *E. coli* for 1h and IL-1β measured in peritoneal lavage and serum by ELISA. n = 3 per group, two-tailed t-test. **H**, WT and *Mt3^-/-^* mice primed *i.p.* with poly(I:C) (10 mg/kg) for 6h and challenged with LPS (2 mg/kg) *i.p.* After 18h, IL-1β was measured in peritoneal lavage and serum by ELISA, n = 3 / group, two-tailed t-test. **I**, Bacterial growth in WT and *Mt3^-/-^* mice infected *i.n.* with *K. pneumoniae* (4 X10^4^ CFUs/mouse) for 48h, n = 8-12 per group, two-tailed t-test, data are mean ± SEM.

### MT3 dampens antibacterial resistance and caspase-11 inflammasome activation *in vivo*

We examined whether MT3 regulated antibacterial immunity and non-canonical inflammasome activation *in vivo*. WT and *Mt3^-/-^* mice were infected with *E. coli* intraperitoneally (*i.p.*) for 6h. Compared to WT mice, MT3 deficiency bolstered bacterial elimination from the blood and moderately improved bacterial clearance in the kidney and peritoneal lavage **(Fig. 2E)**. Caspase-11, GSDMD and caspase-1 activation were heightened in kidney homogenates of infected *Mt3^-/-^* mice compared to infected WT mice **(Fig. 2F)**. The decrease in pro-GSDMD of *Mt3^-/-^* mice may be explained by increased conversion of pro- to active-GSDMD form **(Fig. 2F)**. IL-1β in the peritoneal lavage was significantly elevated (p<0.01) and serum IL-1β exhibited a trend towards increase in infected *Mt3^-/-^* mice compared to WT controls **(Fig. 2G)**. To determine if this response is consistent upon LPS challenge *in vivo*, we primed mice *i.p.* with poly(I:C) for 6h and challenged them *i.p.* with LPS. After 18h, IL-1β was elevated in the peritoneal lavage and serum of *Mt3^-/-^* mice compared to WT mice **(Fig. 2H)**.

We then investigated whether MT3 increased susceptibility to other gram-negative bacteria. WT and *Mt3^-/-^* mice were infected intranasally (*i.n.*) with a virulent, heavily encapsulated strain of *Klebsiella pneumoniae* (KP2 2-70). Mice lacking MT3 exhibited improved clearance of *K. pneumoniae* in the spleen, lung and kidney compared to WT mice **(Fig. 2I)**. Gram-positive bacteria activate caspase-11 via the NLRP6 inflammasome ^24^. We determined whether the increased non-canonical inflammasome activation and antibacterial resistance observed in *Mt3^-/-^* mice extended to gram-positive bacterial infection. WT and *Mt3^-/-^* mice were challenged subcutaneously (*s.q.*) with a clinical isolate of Group-A-Streptococcus GAS5448 ^25^. After 72h, *Mt3^-/-^* mice manifested significantly (p<0.01) reduced GAS burden in the kidney and spleen compared to WT mice. Bacterial CFUs in the blood exhibited a similar trend **(Extended Data Fig. 2A)**. Importantly, although *Mt3^-/-^* mice exhibited higher activation of caspase-1 and IL-1β, the levels of active caspase-11 and pro-caspase-11 were diminished in GAS-infected *Mt3^-/-^* mice compared to WT mice **(Extended Data Fig. 2B)**. Thus, although MT3 compromises resistance to both gram-negative and gram-positive bacteria, it specifically suppresses non-canonical inflammasome signaling in response to gram-negative microbial triggers.

### Caspase-11 synergizes with MT3 in impairing *E. coli* clearance *in vivo*

Caspase-11 is crucial in antibacterial defenses particularly against gram-negative bacteria, although some studies have suggested a detrimental role for caspase-11 in bacterial elimination ^26,27,28,29,30,31^. MT3 suppressed antibacterial immunity as well as non-canonical inflammasome activation. Thus, we investigated whether the heightened immunity to *E. coli* in *Mt3^-/-^* mice was due to increased non-canonical inflammasome activation. We infected WT, *Casp-11^-/-^*, *Mt3^-/-^*, and *Mt3^-/-^Casp-11^-/-^* mice **(Extended Data Fig. 3A)** *i.p.* with *E. coli* and assessed bacterial burden 6h post-infection. Compared to WT mice, bacterial elimination was enhanced in *Mt3^-/-^* and *Casp-11^-/-^* mice but this response was further exacerbated in *Mt3^-/-^* mice lacking caspase-11 **(Fig. 3A)**. These data indicate that the combined absence of MT3 and caspase-11 improves resistance to gram-negative bacterial infection. We then analyzed caspase-11 inflammasome mediators in kidney homogenates harvested 6h post-infection. Mice lacking MT3, or caspase-11, exhibited elevated activation of GSDMD, caspase-1 and IL-1β compared to WT mice **(Fig. 3B)**.

**Fig. 3.**
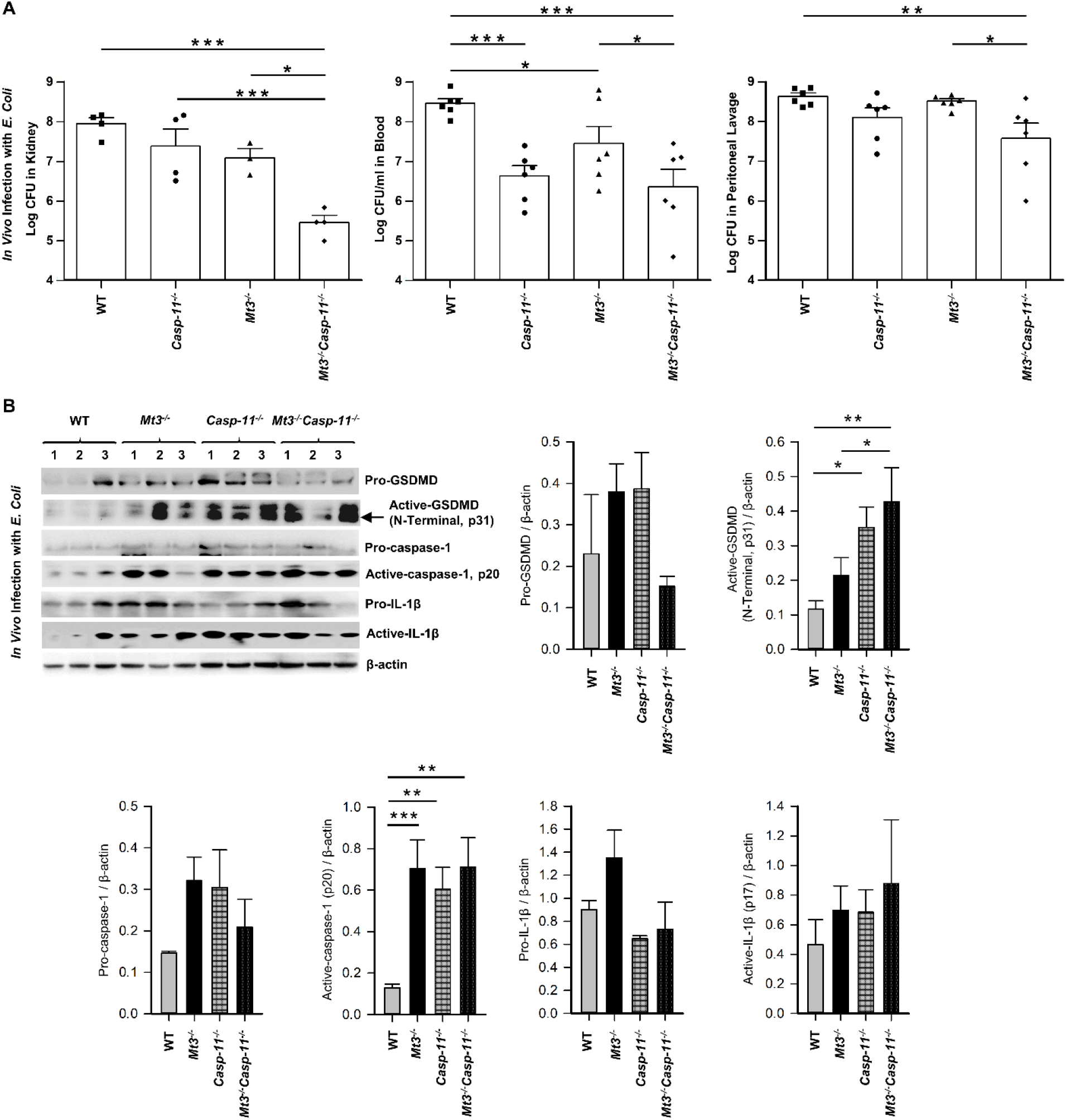
Caspase-11 synergizes with MT3 in impairing bacterial clearance. **See also Extended Data Fig. 3 |** WT, C*asp-11^-/-^*, *Mt3^-/-^* and *Casp-11^-/-^Mt3^-/-^* mice were infected *i.p.* with *E. coli* (1 X10^9^ CFUs/mouse) for 6h. **A**, Bacterial CFUs measured in kidney, blood and peritoneal lavage, n = 3-6 per group, one-way ANOVA. **B**, Western blots of pro- GSDMD, active-GSDMD (p31), pro-caspase-1, active-caspase-1, pro-IL1β and active-IL- 1β in kidney homogenates, n = 3-6 per group, one-way ANOVA, data are mean ± SEM.

These changes were also observed in the *Mt3^-/-^Casp-11^-/-^* mice. Since GSDMD is a target of caspase-11 as well as caspase-1 ^32^, an elevation in active GSDMD may result from higher caspase-1 activation observed in mice lacking MT3, caspase-11 or both. Caspase- 8, a pro-apoptotic caspase, collaborates with caspase-11 to mediate systemic inflammation and septic shock ^33, 34^. Moreover, caspase-8, in addition to caspase-1 can directly cleave IL-1β. We therefore analyzed caspase-8 in kidney homogenates from *E. coli* infected WT, *Mt3^-/-^*, *Casp-11^-/-^* and *Mt3^-/-^Casp-11^-/-^* mice. Caspase-11 negatively influenced the activation of caspase-8 **(Extended Data Fig. 3B)**. These data demonstrate that MT3 negatively controls activation of the non-canonical inflammasome and that both MT3 and caspase-11 cripple resistance to bacterial infection.

### Myeloid MT3 orchestrates negative control of the non-canonical inflammasome

To affirm that the effects on caspase-11 inflammasome activation observed in the *Mt3^-/-^* mice were dependent on myeloid-MT3, we generated mice specifically lacking MT3 in myeloid cells (*Lys2Cre Mt3^fl/fl^*) **(Fig. 4A and Extended Data Fig. 4)**. Genotyping analysis of BMDMϕ and peritoneal Mϕ (PMϕ) from *Lys2Cre*, *Mt3^fl/fl^* and *Lys2Cre Mt3^fl/fl^* mice demonstrated efficient removal of the *Mt3* gene from BMDMϕ and PMϕ only in *Lys2Cre Mt3^fl/fl^* mice **(Fig. 4B)**. To determine if myeloid MT3 deficiency augmented non-canonical inflammasome activation *in vivo*, we infected *Lys2Cre* and *Lys2Cre Mt3^fl/fl^* mice *i.p.* with *E. coli*. After 6h, caspase-11 inflammasome targets and bacterial burden were examined in kidney and blood. *Lys2Cre Mt3^fl/fl^* mice exerted increased activation of caspase-11, GSDMD, caspase-1, and IL-1β in kidney homogenates and improved bacterial elimination compared to *Lys2Cre* control mice **(Figs. 4C and 4D)**. Thus, myeloid MT3 facilitates subversion of non-canonical inflammasome activation and contributes to antibacterial immunity *in vivo*.

**Fig 4.**
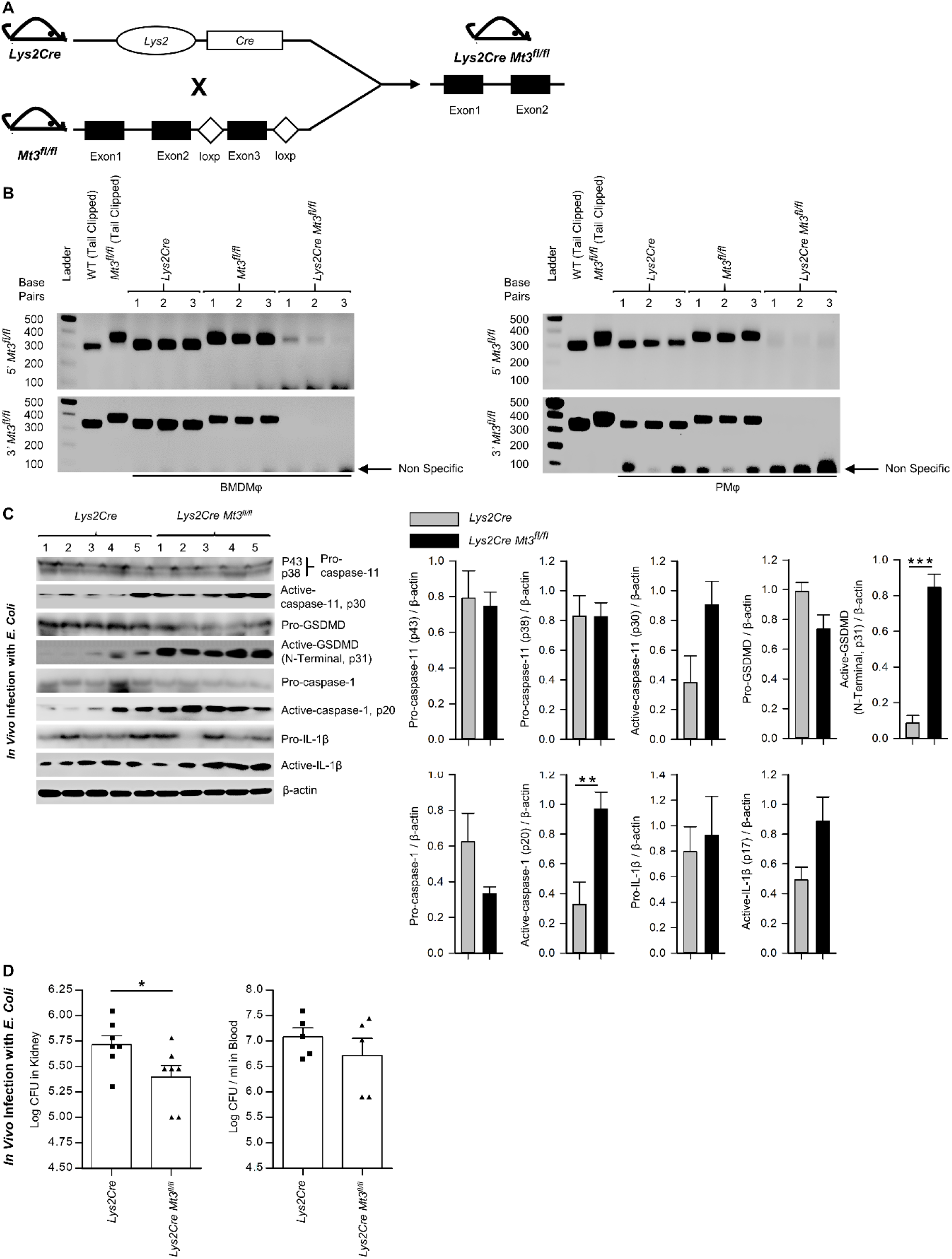
| Myeloid-MT3 suppresses non-canonical inflammasome activation and blunts gram-negative bacterial clearance *in vivo*. See also Extended Data Fig. 4 | A, Generation of *Mt3^fl/fl^* mice by inserting loxp sites flanking exon 3 of the *Mt3* gene using the CRISPR-Cas9 gene targeting approach. *Mt3^fl/fl^* mice crossed with *Lys2Cre* mice to obtain *Lys2Cre Mt3^fl/fl^* mice. **B**, Efficacy of myeloid *Mt3* deletion assessed by genotyping peritoneal Mϕ (PMϕ) and BMDMϕ from *Lys2Cre*, *Mt3^fl/fl^* and *Lys2Cre Mt3^fl/fl^* mice. Gel electrophoresis analysis demonstrating efficient deletion of the *Mt3* gene from BMDMϕ and PMϕ of *Lys2Cre Mt3^fl/fl^* mice. **C**, Western blots of pro-caspase-11, active-caspase- 11, pro-GSDMD, active-GSDMD (p31), pro-caspase-1, active-caspase-1, pro-IL1β and active-IL-1β in whole kidney homogenates of mice infected as above, n = 3-5 per group, two-tailed t-test. **D**, Bacterial CFUs in kidney and whole blood of *Lys2Cre* and *Lys2Cre Mt3^fl/fl^* mice infected *i.p.* with *E. coli* (1 X10^9^ CFUs/mouse) for 6h, n = 3-5 per group, two- tailed t-test, data are mean ± SEM.

### MT3 exerts a brake on the TRIF-IRF3-STAT1 axis to curtail caspase-11 signaling

Signaling via the TRIF pathway is crucial for caspase-11 activation and synergistic engagement of the NLRP3 inflammasome leading to activation of caspase-1 and IL-1β ^20^. Downstream of TRIF, IRF3 and IRF7 induce IFNβ production that activates STAT1 signaling and promotes transcription of inflammasome components including caspase-11 and guanylate binding proteins (GBPs). GBP2 and GBP5 facilitate LPS release into the cytosol from intracellular vacuoles containing bacteria ^35, 36^. Our functional enrichment data based on protein-protein interaction network analyses revealed a potential involvement of MT3 in LPS, TRIF, type-I IFN and IL-1 signaling **(Table 1 and Extended Data Table 1, Extended Data Files 1 and 2)**. We further examined our published RNA-seq data (NCBI SRA: PRJNA533616) to determine differentially expressed genes by comparing resting WT and *Mt3^-/-^* BMDMϕ ^19^. The derived list of differentially expressed genes significantly enriched 12 GO BP categories directly related to cytokine and chemokine signaling and regulation of inflammatory responses based on DAVID functional enrichment analysis **(Fig. 5A)** ^37^. These analyses suggested that MT3 deficiency perturbed the expression of immune-related genes even at the resting state. We reported that a lack of MT3 augments IFNγ responsiveness ^19^. Herein, from our RNA-seq analysis, we identified 20 genes related to IFN-signaling that were upregulated in resting *Mt3^-/-^* BMDMϕ compared to resting WT BMDMϕ (p adj <0.05) **(Figs. 5B and 5C)**. Among these, *Isg15*, *Mx1* and *Ifit* (*Ifit1bl1*, *Ifit3*, *Ifit3b*, *Ifit2*, *Ifit1*, *Ifit1bl2*) family of genes are known targets of type-I IFNs ^38,39,40,41^. These observations led us to posit that MT3 regulated the cellular response to LPS challenge by modulating the TRIF-IRF3-STAT1 axis upstream of non-canonical inflammasome activation. LPS engages the TRIF-IRF3- STAT1 axis via toll-like receptor 4 (TLR4) signaling in Mϕ. To address this hypothesis, we challenged WT and *Mt3^-/-^* BMDMϕ with iLPS or vehicle and examined activation of the TRIF-IRF3-STAT1 pathway. Mϕ lacking MT3 exerted increased activation of phospho- IRF3 (pIRF3), pSTAT1, GBP2 and GBP5 **(Fig. 5D)**. TRIF protein levels was unaltered by MT3 deficiency **(Fig. 5E)**. Type-I IFN signaling is required for activation of the caspase- 11 inflammasome cascade by gram-negative bacteria ^20^. As MT3 deficiency augmented the expression of genes involved in IFN signaling **(Figs. 5B and 5C)**, we blocked the interferon-α/β receptor (IFNAR)1 using a monoclonal antibody prior to iLPS challenge in WT and *Mt3^-/-^* Mϕ. IFNAR1 blockade resulted in decreased pro-caspase-11 (p43 subunit). Total STAT1 and pro-caspase-1 (p38 subunit) were not greatly affected **(Extended Data Fig. 5)**. We found robust attenuation of pSTAT1, active-caspase-11 and active-caspase- 1, but secretion of active-IL-1β in both WT and *Mt3^-/-^* Mϕ was increased upon blockade of IFNAR1 signaling. This finding corresponded with high pro-IL -1β levels in Mϕ treated with the IFNAR1 antibody **(Extended Data Fig. 5)**. These data indicate that although IFNAR1 signaling is required for fueling the non-canonical inflammasome cascade and activation of caspase-1, pro-IL-1β and its activation are suppressed by IFNAR1.

**Fig. 5.**
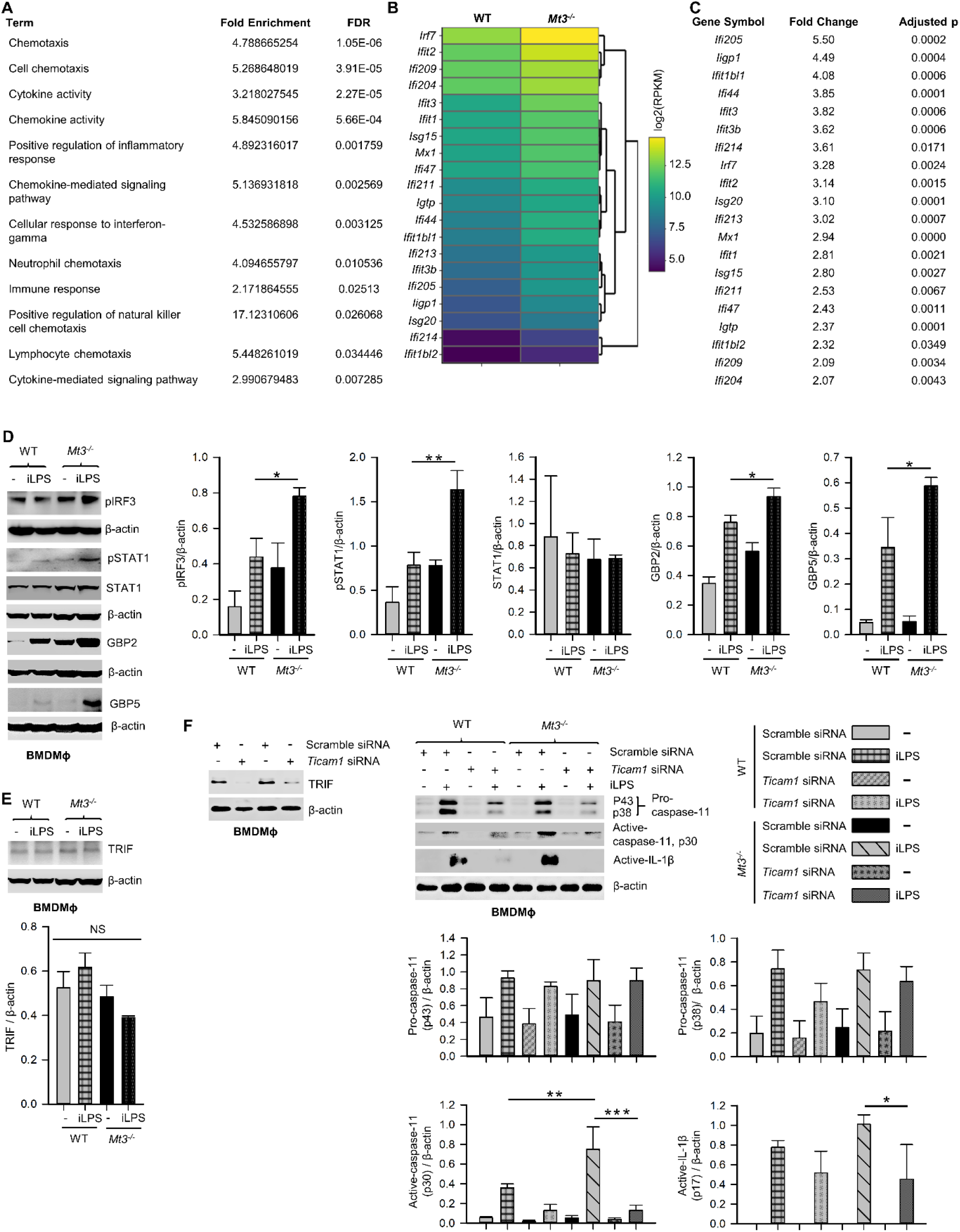
MT3 thwarts TRIF-IRF3-STAT1 signaling to suppress non-canonical inflammasome activation. **See also Extended Data Fig. 5 | A**, Functional enrichment analysis of differentially expressed genes using RNA-seq data from resting WT and *Mt3^-/-^* BMDMϕ (NCBI SRA: PRJNA533616) ^19^ FDR, false detection rates. **B**,**C**, Heat map (left) and table (right) show differentially expressed IFN-related genes in resting *Mt3^-/-^* BMDMϕ compared to resting WT BMDMϕ obtained from RNA-seq analysis. **D**, Western blots of pIRF3, pSTAT1, STAT1, GBP2 and GBP5 in vehicle or iLPS (10 μg/ml)-treated WT and *Mt3^-/-^* BMDMϕ lysates, 3-4 independent experiments, one-way ANOVA. **E**, Western blots of TRIF in lysates from WT and *Mt3^-/-^* BMDMϕ stimulated as above, 3 independent experiments, one-way ANOVA. **F**, Scramble and *Ticam1* siRNA treated WT and *Mt3^-/-^* BMDMϕ treated with iLPS (10 μg/ml) or vehicle for 48h. Immunoblots of TRIF, pro- caspase-11, and active-caspase-11 in lysates and active-IL-1β in supernatants, 2 independent experiments, 3 independent experiments, one-way ANOVA, data are mean ± SEM.

We queried if MT3 exerted a brake on TRIF signaling to downmodulate non-canonical inflammasome activation. WT and *Mt3^-/-^* BMDMϕ were treated with scramble or *Ticam1* (gene encoding TRIF) siRNA, and challenged with iLPS **(Fig. 5F)**. *Ticam1* silencing reversed the effects of MT3 deficiency resulting in a sharp reduction in caspase- 11 and IL-1β activation in Mϕ **(Fig. 5F)**. These data reveal a central role for MT3 in attenuating the crosstalk between TRIF signaling and the caspase-11 activation cascade.

### Zn^2+^ flux by MT3 drives suppression of the non-canonical inflammasome in Mϕ

MTs are master regulators of intracellular Zn^2+^ availability and distribution ^42, 43^. We determined if negative control of the non-canonical inflammasome by MT3 was Zn^2+^- dependent. First, we systematically assessed Zn^2+^ changes in WT and *Mt3^-/-^* BMDMϕ upon challenge with iLPS over time using SEC-ICP-MS. Activation of the non-canonical inflammasome in WT Mϕ was associated with profound changes in the intracellular Zn^2+^ pool. iLPS exposure led to a gradual increase in total Zn^2+^ largely associated with the chromatogram peak(s) between 18-21 min. that we previously identified as MTs **(Figs. 6A and 6B)** ^9, 11^. The time-dependent elevation in Zn^2+^ corresponded with kinetics of *Mt3* induction **(Figs. 6A, 6B and 1A)**. Mϕ lacking MT3 failed to elevate total Zn^2+^ and MT- associated Zn^2+^ in response to iLPS **(Fig. 6B)**. In contrast to the increase in Zn^2+^ pool observed in WT Mϕ challenged with iLPS, resting *Mt3^-/-^* Mϕ harbored higher Zn^2+^ content that reduced over time post iLPS challenge **(Fig. 6B)**. These data indicate that MT3 drives an elevation in intracellular Zn^2+^ in Mϕ during non-canonical inflammasome activation.

**Fig. 6.**
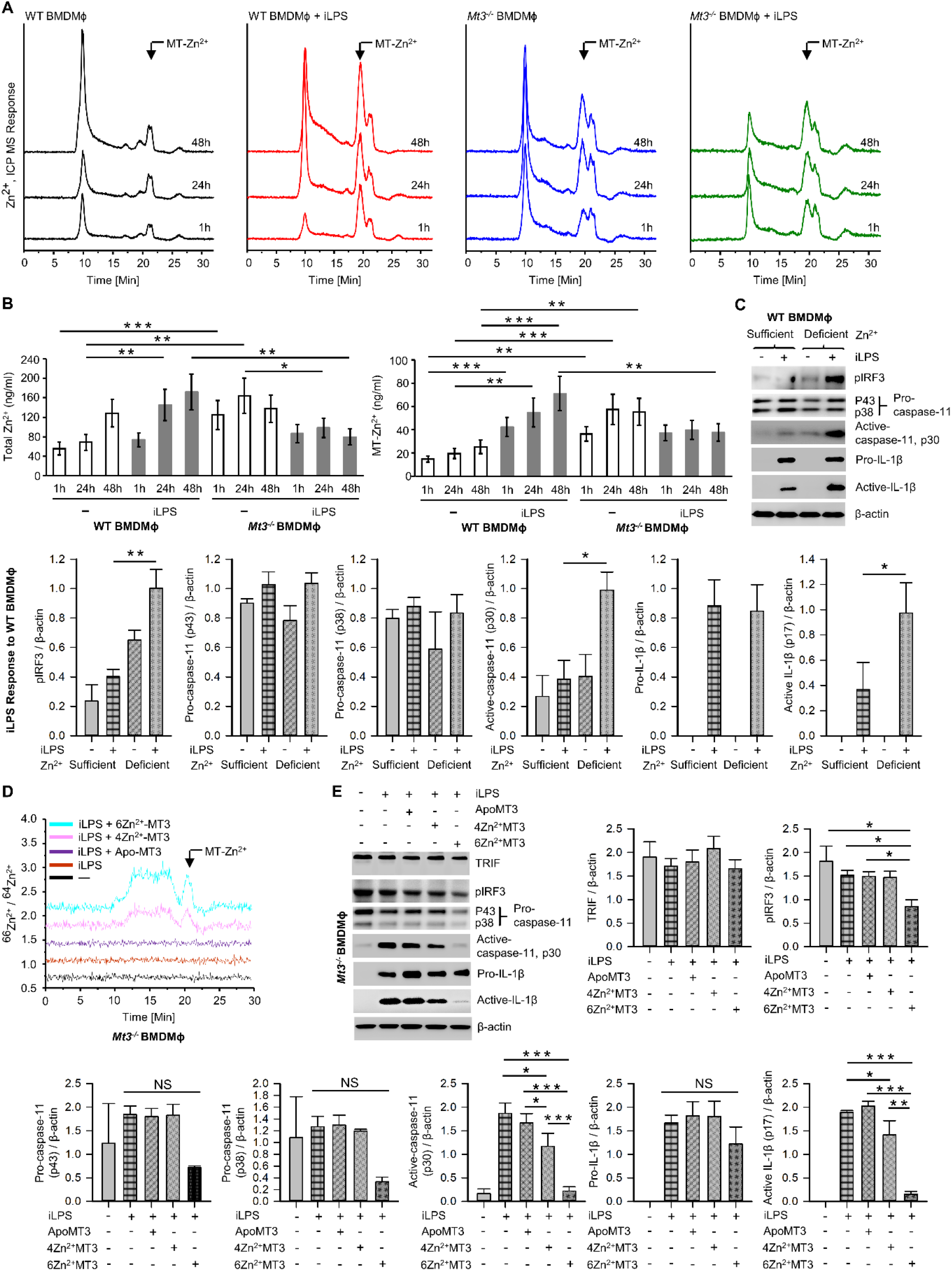
MT3-Zn^2+^ axis drives negative regulation of the non-canonical inflammasome. **See also Extended Data Figs. 6 and 7 | A**, SEC-ICP-MS of WT and *Mt3^-/-^* BMDMϕ exposed to vehicle or iLPS (10 ug/ml) for the indicated time points, chromatograms depict Zn^2+^ distribution in cell lysates across various molecular masses, arrow indicates Zn^2+^ associated with the MT-peak (18-21 min.) on the chromatogram, Y axis is off-set to allow easy comparison under the same scale. **B**, Bar graphs of total Zn^2+^ and MT-Zn^2+^ in WT and *Mt3^-/-^* BMDMϕ post iLPS (10 ug/ml) or vehicle exposure. Two-way t-test against respective BMDMϕ controls at each time point, 3 independent experiments, data are mean ± SD. **C**, WT BMDMϕ treated with iLPS (10 μg/ml) or vehicle for 24h in Zn^2+^ sufficient or Zn^2+^ deficient Opti-MEM media, immunoblots of pIRF3, pro-caspase-11, active-caspase-11 and pro-IL-1β in lysates and active-IL-1β in media supernatants, one-way ANOVA, data are mean ± SEM. **D**, *Mt3^-/-^* BMDMϕ transfected with Pro-Ject^TM^ or Pro-Ject^TM^ complexed with apo-MT3, 4Zn^2+^MT3 or 6Zn^2+^MT3 and treated with iLPS (10 μg/ml) or vehicle for 24h in Zn^2+^ deficient Opti-MEM media. Chromatograms depict Zn^2+^ distribution in cell lysates across various molecular masses, arrow indicates Zn^2+^ signal associated with the MT-peak (18-21 min.) on the chromatogram, Y axis is off- set to allow easy comparison under the same scale. **E.** *Mt3^-/-^* BMDMϕ transfected MT3- Zn^2+^ complexes and treated with iLPS as above, western blots of pIRF3, pro-caspase-11, active-caspase-11 and pro-IL1β in lysates and active-IL-1β in supernatants, one-way ANOVA, data are mean ± SEM.

Zn^2+^ chelation in human monocytes increases IRF3 activation ^44^. We reasoned that if the effects of MT3 were Zn^2+^ dependent, altering the intracellular Zn^2+^ concentration will at least in part reverse the heightened non-canonical inflammasome signaling observed in *Mt3^-/-^* cells. To test this postulate, we exposed WT and *Mt3^-/-^* BMDMϕ to increasing amounts of ZnSO4 and challenged them with iLPS *in vitro*. Exogenous ZnSO4 supplementation remarkably reduced the ability of *Mt3^-/-^* Mϕ to respond to iLPS. pIRF3, pSTAT1 and activation of caspase-11 were reduced in ZnSO4-supplemented *Mt3^-/-^* Mϕ **(Extended Data Fig. 6A)**. A similar effect of Zn^2+^ was also observed in WT Mϕ **(Extended Data Fig. 6A)**. We investigated if exposing WT Mϕ to a Zn^2+^-deficient environment would mimic the effects MT3 deficiency on the non-canonical inflammasome. WT BMDMϕ were cultured in Zn^2+^-sufficient or Zn^2+^-deficient Opti-MEM media prior to iLPS exposure. WT Mϕ exposed to a Zn^2+^-deficient milieu manifested higher pIRF3 and caspase-11 activation accompanied by increased activation and release of IL-1β **(Fig. 6C)**. The amount of TRIF, pro-caspase-11 and pro-IL-1β were not affected by Zn^2+^ deficiency **(Fig. 6C and Extended Data Fig. 6B)**.

Next, we directly addressed whether Zn^2+^ is required for the suppressive function of MT3 on the caspase-11 inflammasome. We first overexpressed the MT3 in *Mt1^-/-^Mt2^-/-^* Mϕ and isolated the protein. *Mt1^-/-^Mt2^-/-^* BMDMϕ were transfected with the *Mt3* overexpressing vector (pCMV6-Ac-MT3-GFP) or an empty vector (pCMV6-Ac-GFP) control. *Mt1^-/-^Mt2^-/-^* Mϕ were used so as to exclude any contribution of these MTs in the MT3 purification process. The MT-associated peak from MT3-overexpressed Mϕ was identified by SEC-ICP-MS **(Extended Data Fig. 7)** and collected. We complexed MT3 with the ^66^Zn^2+^ isotope to acquire an MT3-Zn^2+^ saturation of 4 Zn^2+^ ions per MT3 (MT3- 4Zn^2+^) and 6 Zn^2+^ ions per MT3 (MT3-6Zn^2+^). The ^66^Zn^2+^ isotope was used to monitor changes in the ratio of ^66^Zn^2+^ / ^64^Zn^2+^ post-transfection of the MT3-^66^Zn^2+^ complexes in Mϕ. We transfected apo-MT3, MT3-4Zn^2+^ or MT3-6Zn^2+^ into *Mt3^-/-^* BMDMϕ in Zn^2+^- deficient media. The use of *Mt3^-/-^* BMDMϕ and Zn^2+^-deficient media enabled exclusion of any possible contribution from endogenous MT3 and exogenous Zn^2+^ in our analysis. Post-transfection of apo-MT3 or MT3-Zn^2+^ complexes, Mϕ were challenged with iLPS to activate the non-canonical inflammasome. To confirm that intracellular Zn^2+^ changes occurred upon transfection of the MT3- ^66^Zn^2+^ complexes, we analyzed BMDMϕ lysates by SEC-ICP-MS. *Mt3^-/-^* cells transfected with MT3-4Zn^2+^ and MT3-6Zn^2+^ but not apo-MT3 exhibited an increase in the ^66^Zn^2+^ / ^64^Zn^2+^ ratio in the MT-peak region at 18-21 mins. in the chromatogram indicating an elevation in the intracellular ^66^Zn^2+^ isotope **(Fig. 6D)**. These data confirm that transfection of the MT3-Zn^2+^ complexes resulted in an increase in intracellular ^66^Zn^2+^ in Mϕ. In parallel, we isolated cell lysates and supernatants proteins from these Mϕ to determine whether apo-MT3 or the MT3-Zn^2+^ complexes modulated the non-canonical inflammasome pathway. Transfection of MT3-4Zn^2+^ and MT3-6Zn^2+^ but not apo-MT3, dampened pIRF3, active caspase-11 and active IL-1β in response to iLPS **(Fig. 6E)**. TRIF levels were unaffected by MT3 transfection **(Fig. 6E)**. Pro-caspase-11 and pro-IL-1β were modestly diminished by MT3-4Zn^2+^ and MT3-6Zn^2+^ exposure, but these changes were not significant **(Fig. 6E)**. The effect of MT3-6Zn^2+^ was more profound than that of MT3-4Zn^2+^ indicating that a higher Zn^2+^ saturation on MT3 corresponded with a stronger suppressive effect on the non-canonical inflammasome **(Fig. 6E)**.

Taken together, these findings reveal a previously undescribed interplay between the non-canonical inflammasome and its negative regulator, whereby the MT3-Zn^2+^ axis suppresses caspase-11 inflammasome, but the two molecules concur in compromising immunological fitness of the host during bacterial pathogenesis.

## Discussion

Human CASPASE-4 or mouse caspase-11 are inflammatory caspases that drive cell death via pyroptosis. These caspases directly recognize bacterial LPS in the cytosol resulting in CASPASE-4 or caspase-11 auto-processing and synergistic activation of the NLRP3 inflammasome that culminates in caspase-1 activation, processing and release of IL-1β and IL-18 ^2, 28^. While the non-canonical inflammasome boosts host immunological fitness to some bacterial infections, heightened activation of this cascade poses the danger of tissue injury and organ failure. Thus far, negative regulation of IFNβ production by prostaglandin E2, immunity-related GTPases M clade, cyclic-adenosine monophosphate, and low dose oxidized phospholipid oxPAPC have been shown to thwart activation of the non-canonical inflammasome ^45,46,47,48^. The role of MTs in regulating inflammasome activation pathways has largely been unknown. Herein, we identify a previously undescribed function of MT3 in curtailing the highly inflammatory non-canonical inflammasome activation cascade via Zn^2+^ regulation. We demonstrate that while MT3 orchestrates negative regulation of the caspase-11 inflammasome, the combined presence of MT3 and caspase-11 blunts resistance to *E. coli* infection *in vivo*. These studies illuminate a central role for the MT3-Zn^2+^ axis in shaping the intricate balance between host antibacterial immunity and unrestrained inflammation.

MT1 and MT2 are ubiquitously expressed and can be induced by infection ^49,50,51,52^. Initial studies on MT3 revealed tissue-restricted expression with high levels predominantly found in the brain tissue where it inhibits neuronal cell death ^21, 53^. The immunological functions of MT3, particularly in the innate arm have only recently been investigated ^11, 19, 21^. We reported that *Mt3* is inducible by the pro-resolving cytokines IL- 4 and IL-13 in Mϕ. One inducer of *Mt3* expression is STAT6 signaling, and this MT is crucial in shaping the phenotypic and metabolic attributes of Mϕ stimulated with type-2 cytokines ^11, 19^. Studies on MTs in response to exogenous LPS stimulation have largely focused on MT1 and MT2. Monocytes and Mϕ induce MT1 and MT2 upon extracellular LPS exposure ^54, 55^. We found that iLPS challenge also induced *Mt1* and *Mt2* in Mϕ, although their expression receded to baseline over time. In contrast, *Mt3* expression gradually increased as non-canonical inflammasome activation progressed. Subversion of inflammatory responses in Mϕ by MT3 and the delayed expression pattern in response to a non-canonical inflammasome trigger are reminiscent of waning inflammation after the initial peak of inflammasome activation has subsided ^19^. In line with this hypothesis, protein interaction network analysis predicted the involvement of MT3 in cellular responses related to non-canonical inflammasome activation. Mϕ lacking MT3 exerted robust activation of caspase-11, caspase-1 and IL-1β. Similar to our observations with mouse MT3, a lack of human MT3 exacerbated the activation of CASPASE4 and IL-1β in hMϕ. Human and mouse MT3 proteins that share 86% identity thus have consistent roles that culminate in negative regulation of the non-canonical inflammasome cascade in Mϕ ^56^. As non-canonical inflammasome activation progressed, MT3 guarded against its unrestrained activation to avert potential inflammatory damage. Together, these observations reveal a pivotal role for MT3 in curtailing the vigor of the caspase-11 activation cascade.

*In vivo*, LPS released from OMV of gram-negative bacteria triggers caspase-11 activation ^2, 3^. Myeloid-MT3 contributed to averting excessive activation of caspase-11 and synergistic activation of the canonical inflammasome in response to gram-negative microbial triggers. Although MT3 compromised antibacterial resistance to *E. coli* and *K. pneumoniae,* this was not due to its suppressive action on the caspase-11 inflammasome *in vivo*. Instead, *Mt3^-/-^* and *Casp-11^-/-^* mice individually manifested improved antibacterial immunity, an effect that was further augmented when *Mt3^-/-^* mice lacked caspase-11. Of note, the activation of caspase-1 and caspase-8 in the absence of MT3 and caspase-11 reveal that canonical inflammasome activation was operational and both caspase-1 and caspase-8 may contribute to IL-1β activation *in vivo* ^34^.

Mϕ utilize Zn^2+^ deprivation and Zn^2+^ intoxication mechanisms as strategies for antimicrobial defense ^52, 57, 58^. We previously showed that ablation of MT3 in Mϕ augments immunity to *Histoplasma capsulatum* as well as *E. coli*. The increased antimicrobial resistance in *Mt3^-/-^* Mϕ is at least partially attributable to a decrease in the Mϕ exchangeable Zn^2+^ pool and exaggerated IFNγ responsiveness ^11, 19^. The finding that MT3 deficiency dually bolstered inflammasome activation and antibacterial immunity underpins a role for this protein in suppressing the emergence of a proinflammatory phenotype in Mϕ. Our data unveil a unique crosstalk between caspase-11 and its negative regulator, whereby although MT3 keeps caspase-11 activation under control, the two synergistically compromise host immunological fitness to gram-negative bacterial infection.

Caspase-11 activation can have opposing effects on clearance of different bacteria. It improves resistance to *Burkholderia thailandensis*, *B. pseudomallei*, *Brucella abortus*, and *Legionella pneumophila* but may compromise immunity to *B. cenocepacia*, *Salmonella typhimurium*, *E. coli*, *Shigella flexneri*, *K. pneumoniae* and gram-positive infections including *Streptococcus pyogenes*, *Staphylococcus aureus* and *Listeria monocytogenes* ^24, 27,28,29, 31, 59,60,61,62,63,64,65^. Lipoteichoic acid from gram-positive bacteria engages the caspase-11 inflammasome via NLRP6 ^24^. Likewise, GAS infection led to caspase-11 activation *in vivo*. Although MT3 exerted disparate effects on caspase-11 activation in gram-positive and gram-negative infections, caspase-1 and IL-1β activation was suppressed by MT3 in both infection settings. In the context of GAS infection, both the host and the pathogen contribute to canonical inflammasome activation. Surface and secreted GAS virulence factors *emm,* and the streptococcal pyrogenic exotoxin B (SpeB) proteins act as second signals to activate caspase-1 signaling ^66,67,68^. Our data do not exclude the role of pathogen-derived factors in contributing to the increased canonical inflammasome activation observed in *Mt3^-/-^* mice infected with GAS. Nonetheless, our findings indicate that MT3 exerted a suppressive effect on the canonical caspase-1 pathway activated by gram-positive bacteria, while sparing negative regulation of the upstream non-canonical inflammasome activation *in vivo*.

The TRIF pathway is a central node in activation of the caspase-11 inflammasome in response to gram-negative infection ^20^. Targeting TRIF, but not IFNAR1, completely reversed the inflammatory cascade, including IL-1β activation. Blockade of IFNAR1 attenuated downstream activation of STAT1, caspase-11 and caspase-1, but both pro-IL-1β and active IL-1β levels were dramatically enhanced. This finding contrasts with the previously reported requirement of both TRIF and IFNAR1 in this pathway ^20^. Although that study utilized BMDMϕ from *IFNAR1^-/-^* mice and we used an anti-IFNAR1 monoclonal antibody, both approaches resulted in attenuation of targets downstream of IFNAR1. Emerging evidence points to an indirect inhibitory effect of type-I IFNs on inflammasome activation by decreasing pro-IL-1β transcription via IL-10 or 25-hydroxycholesterol ^69, 70^. Interfering with IFNAR1 signaling can therefore subdue activation of critical inflammasome components including caspase-11 and caspase-1 but sustain IL-1β production and activation.

The crucial function of Zn^2+^ as a signaling molecule is well documented ^13, 71,72,73^. Here, we demonstrate that caspase-11 activation is accompanied by a gradual expansion of the intracellular Zn^2+^ pool driven by MT3. Zn^2+^ diminishes signaling via IRF3 by limiting its nuclear localization ^44^. Accordingly, MT3 interfered with signaling via the TRIF-IRF3- STAT1 axis by shaping the Mϕ Zn^2+^ pool. The MT3-Zn^2+^ axis dampened IRF3 phosphorylation and downstream mediators without impacting TRIF levels. Although we cannot rule out the direct effect of Zn^2+^ on inflammasome components downstream of IRF3, the suppressive action of MT3 on the non-canonical inflammasome was Zn^2+^ dependent. Our data demonstrate that by manipulating the Mϕ Zn^2+^ milieu, caspase-11 activation can either be triggered or averted. This finding has important implications in defining a role for Zn^2+^ in subverting caspase-11 driven hyperinflammation during sepsis.

Taken together, our studies illuminate a double-edged phenomenon in inflammasome regulation whereby the MT3-Zn^2+^ axis is a sentinel of the caspase-11 inflammasome but MT3 and the non-canonical inflammasome function in concert to compromise host antibacterial resistance.

## Acknowledgements

We thank Transgenic Animal and Genome Editing Core at Cincinnati Children’s Hospital Medical (CCHMC) Center for production of *Mt3^fl/fl^* mice. This work was supported by a Junior Faculty Pilot Project Award, American Heart Association 19CDA34770022 Award, American Association of Immunology Careers in Immunology Fellowship Awards to K. S. V and NIAIDR01 AI106269-06 awarded in part to K. S. V.

## Author contributions

D. C and K. S. V conducted molecular and biochemical *in vitro* and *in vivo* experiments, generated *Casp-11^-/-^Mt3^-/-^* and *Lys2Cre Mt3^fl/fl^* mice, analyzed data, and wrote the manuscript, J. G conducted *in vivo* infections with *K. pneumoniae*, A. S, S. N and S. M conducted *in vivo* experiments with GAS infection and analyzed data, A. P assisted with bioinformatics analysis, J. L. F conducted chromatographic and mass spectrometric analysis using ICP-MS and SEC-ICP-MS and MT3-Zn^2+^ complex preparations, and analyzed data, K. S. V designed and supervised the project.

## Supplemental materials

**Extended Data Table 1.** Protein-protein interactions network of *Homo sapiens* MT3 with 50 first-shell and 50 second-shell interactors to determine functionally enriched GO BP categories using the STRING database. **See also** Table 1 **and Extended Data Files 1, 2 |**

| Sl. No | Term ID | Term Description | FDR | Protein Labels |
| --- | --- | --- | --- | --- |
| 1 | GO:0060548 | Negative regulation of cell death | 1.12E-12 | Akt1, Ppp5c, Hspb1, Sod2, Hspa8, Gsk3b, Sod1, Hsp90ab1, Hspd1, Cat, Txn1, Dnaja1, Bag1, Cdk5, Bag3, Mt3, Mt1, Ptges3, Bag5, Ar, Pgr, Hsf1, Ahr, Hspa1b |
| 2 | GO:0043067 | Regulation of programmed cell death | 3.01E-11 | Akt1, Hspb1, Sod2, Hspa8, Gsk3b, Sod1, Hsp90ab1, Hspd1, Cat, Ppid, Dnaja1, Bag1, Vcp, Scp2, Cdk5, Bag3, Mt3, Mt1, Tspo, Bag5, Ar, Pgr, Hsf1, Nr3c1, Hdac6, Hspa1b |
| 3 | GO:0042981 | Regulation of apoptotic process | 1.69E-10 | Akt1, Hspb1, Sod2, Hspa8, Gsk3b, Sod1, Hsp90ab1, Hspd1, Cat, Ppid, Dnaja1, Bag1, Vcp, Scp2, Cdk5, Bag3, Mt3, Mt1, Tspo, Bag5, Ar, Pgr, Hsf1, Nr3c1, Hspa1b |
| 4 | GO:0043066 | Negative regulation of apoptotic process | 2.34E-09 | Akt1, Hspb1, Sod2, Hspa8, Gsk3b, Sod1, Hsp90ab1, Hspd1, Cat, Dnaja1, Bag1, Bag3, Mt3, Mt1, Bag5, Ar, Pgr, Hsf1, Hspa1b |
| 5 | GO:0043068 | Positive regulation of programmed cell death | 3.25E-08 | Akt1, Sod2, Gsk3b, Sod1, Hspd1, Ppid, Dnaja1, Bag1, Vcp, Scp2, Cdk5, Tspo, Hsf1, Nr3c1, Hdac6 |
| 6 | GO:2001242 | Regulation of intrinsic apoptotic signaling pathway | 0.0011 | Hspb1, Sod2, Sod1, Dnaja1, Bag5 |
| 7 | GO:0032680 | Regulation of tumor necrosis factor production | 0.0048 | Hspb1, Hspd1, Tspo, Hsf1 |
| 8 | GO:0060334 | Regulation of interferon-gamma-mediated signaling pathway | 0.0059 | Cdc37, Hsp90ab1 |
| 9 | GO:0060338 | Regulation of type I interferon-mediated signaling pathway | 0.0083 | Cdc37, Hsp90ab1 |
| 10 | GO:0032496 | Response to lipopolysaccharide | 0.0107 | Akt1, Sod2, Hspd1, Tspo, Hsf1 |
| 11 | GO:0010508 | Positive regulation of autophagy | 0.016 | Cdc37, Gsk3b, Hdac6 |
| 12 | GO:0002711 | Positive regulation of T cell mediated immunity | 0.0237 | Hspa8, Hspd1 |
| 13 | GO:0001817 | Regulation of cytokine production | 0.0339 | Hspb1, Sod1, Hsp90ab1, Hspd1, Tspo, Hsf1 |
| 14 | GO:0071222 | Cellular response to lipopolysaccharide | 0.0376 | Akt1, Tspo, Hsf1 |
| 15 | GO:0071356 | Cellular response to tumor necrosis factor | 0.0415 | Akt1, Bag4, Cfl1 |

**Extended Data File 1.**
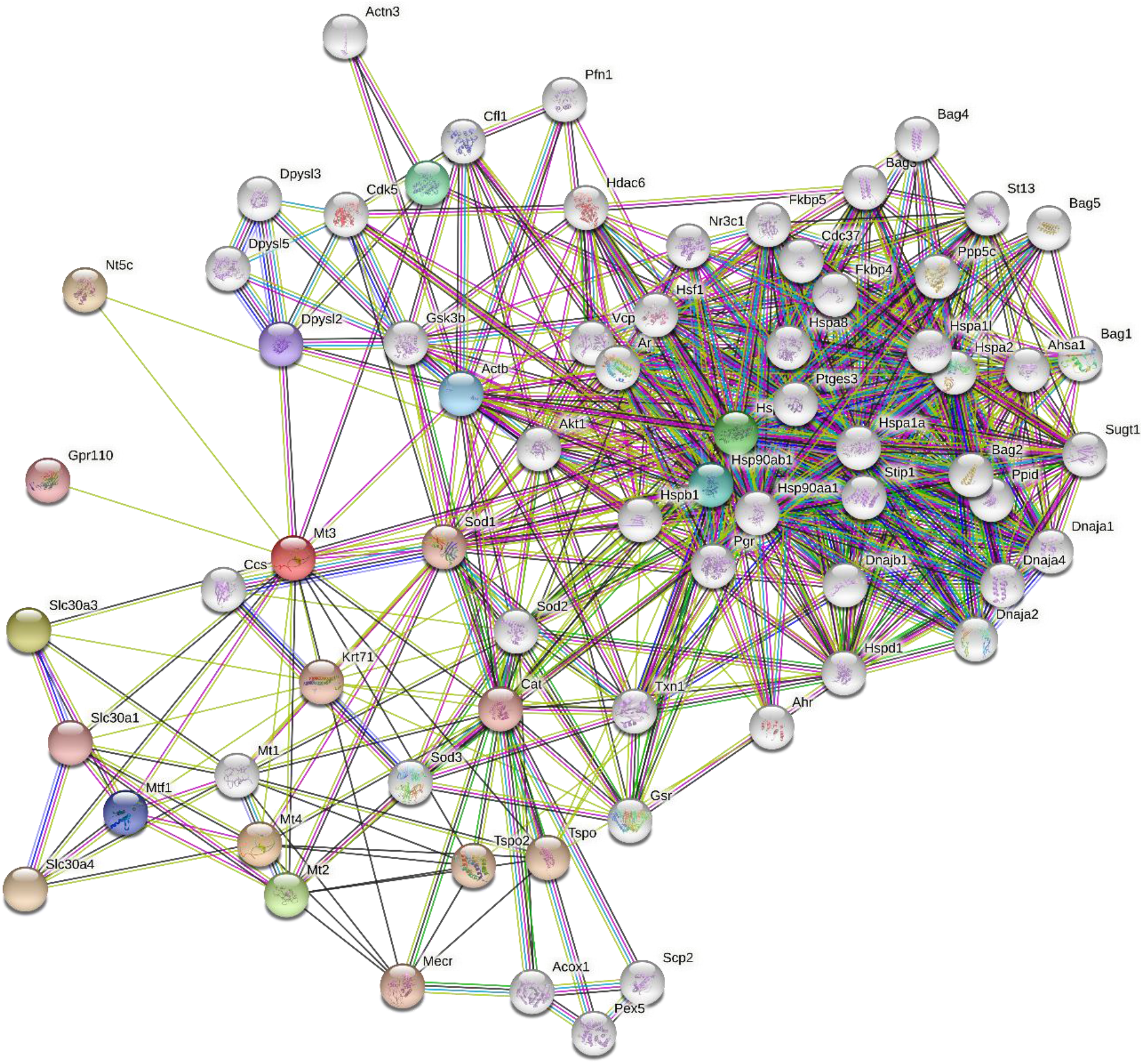

**Extended Data File 2.**
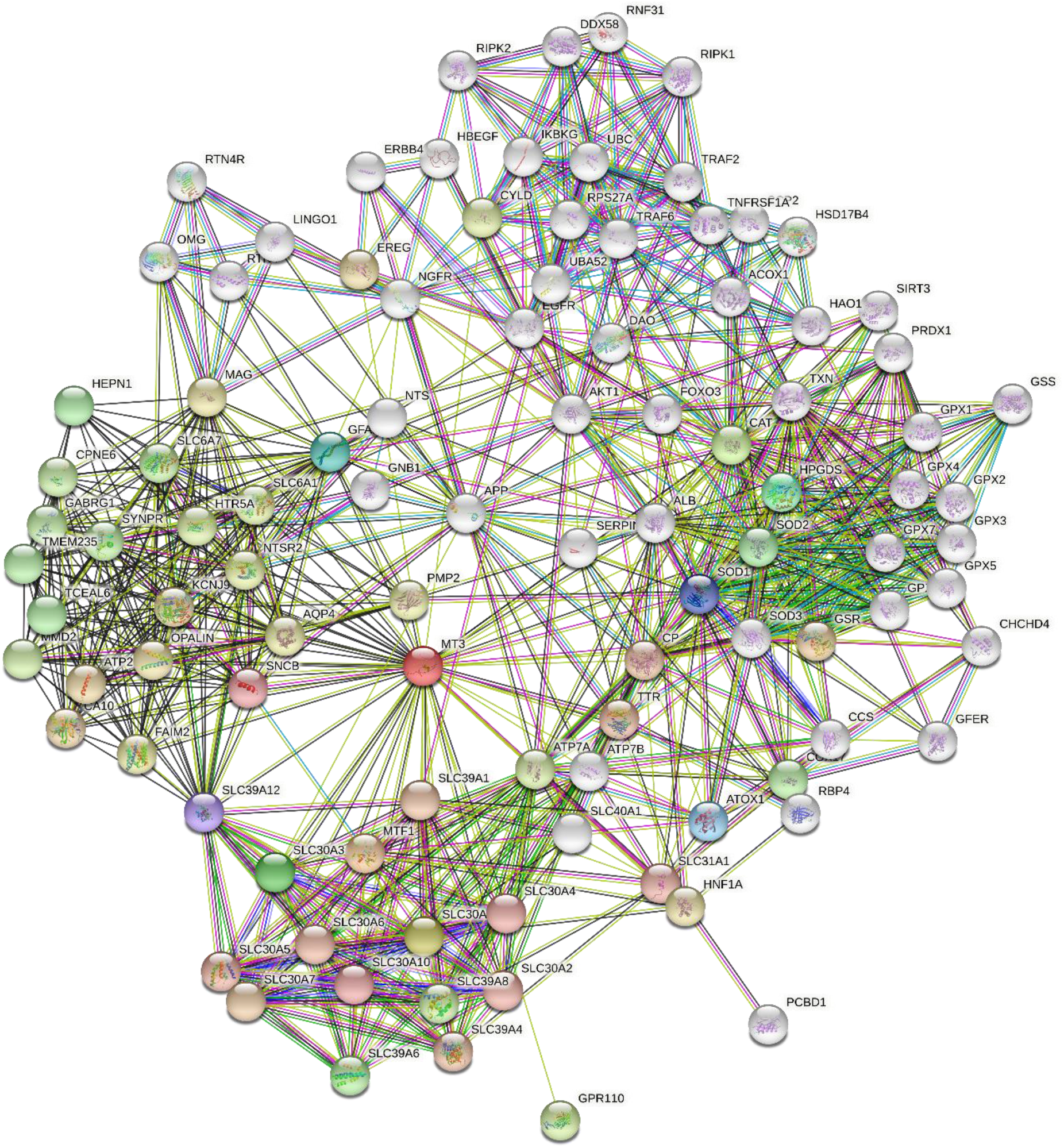
**Extended Data Files 1 and 2. Protein-protein interaction network map of *Homo sapiens* and *Mus musculus* MT3. See also** Table 1 **and Extended** Table 1 **|** Protein-protein interaction network map of MT3 in **(Extended Data File 1)** *Homo sapiens* and **(Extended Data File 2)** *Mus musculus* with 50 first-shell and 50 second-shell interactors showing functionally enriched gene ontology categories for biological processes (GO BP). See table 1 and table S1 for tabular representation of the GO BP categories identified using the STRING database for MT3 protein-protein network interactions in *Mus musculus* and *Homo sapiens*.

**Extended Data Fig. 1.**
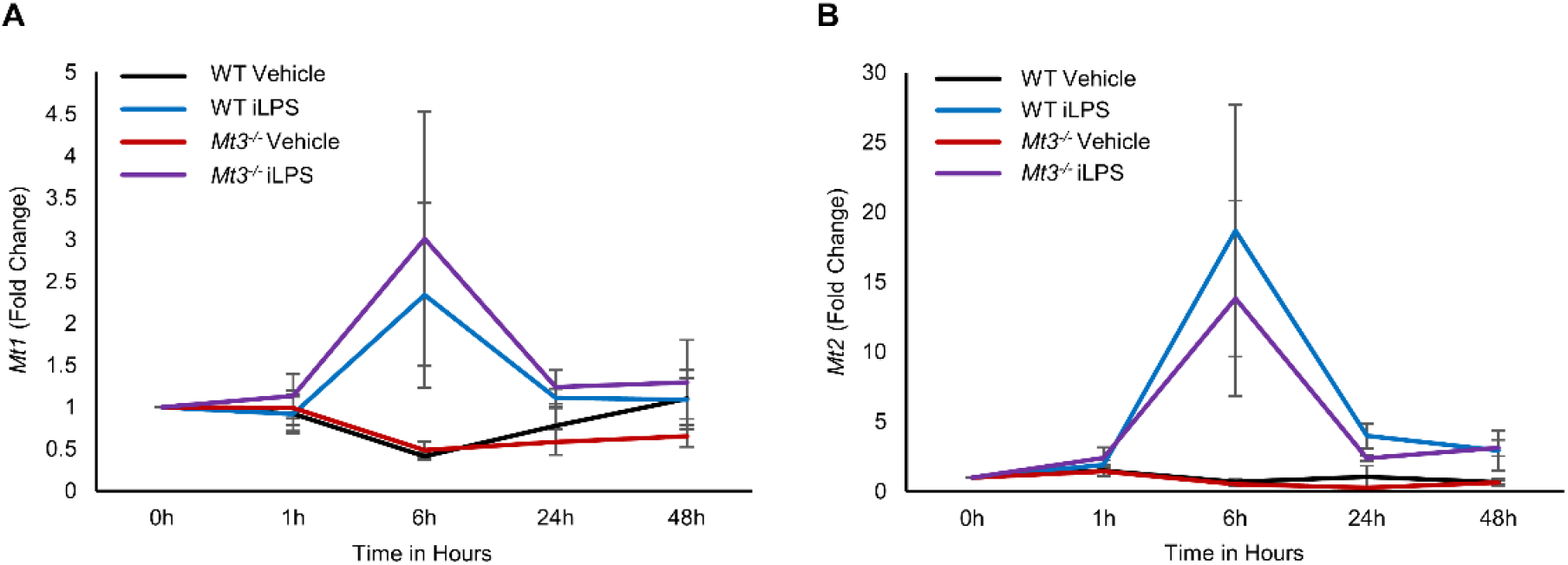
Gene expression of *Mt1* and *Mt2*. **See also Fig. 1 |** qRT-PCR analysis of **(A)** *Mt1* and **(B)** *Mt2* expression over time in WT and *Mt3^-/-^*BMDMϕ stimulated with iLPS (2 μg/ml) or vehicle control. Graphs represent data from 4 independent experiments, t-test (two-tailed p-value), data are mean ± SEM.

**Extended Data Fig. 2.**
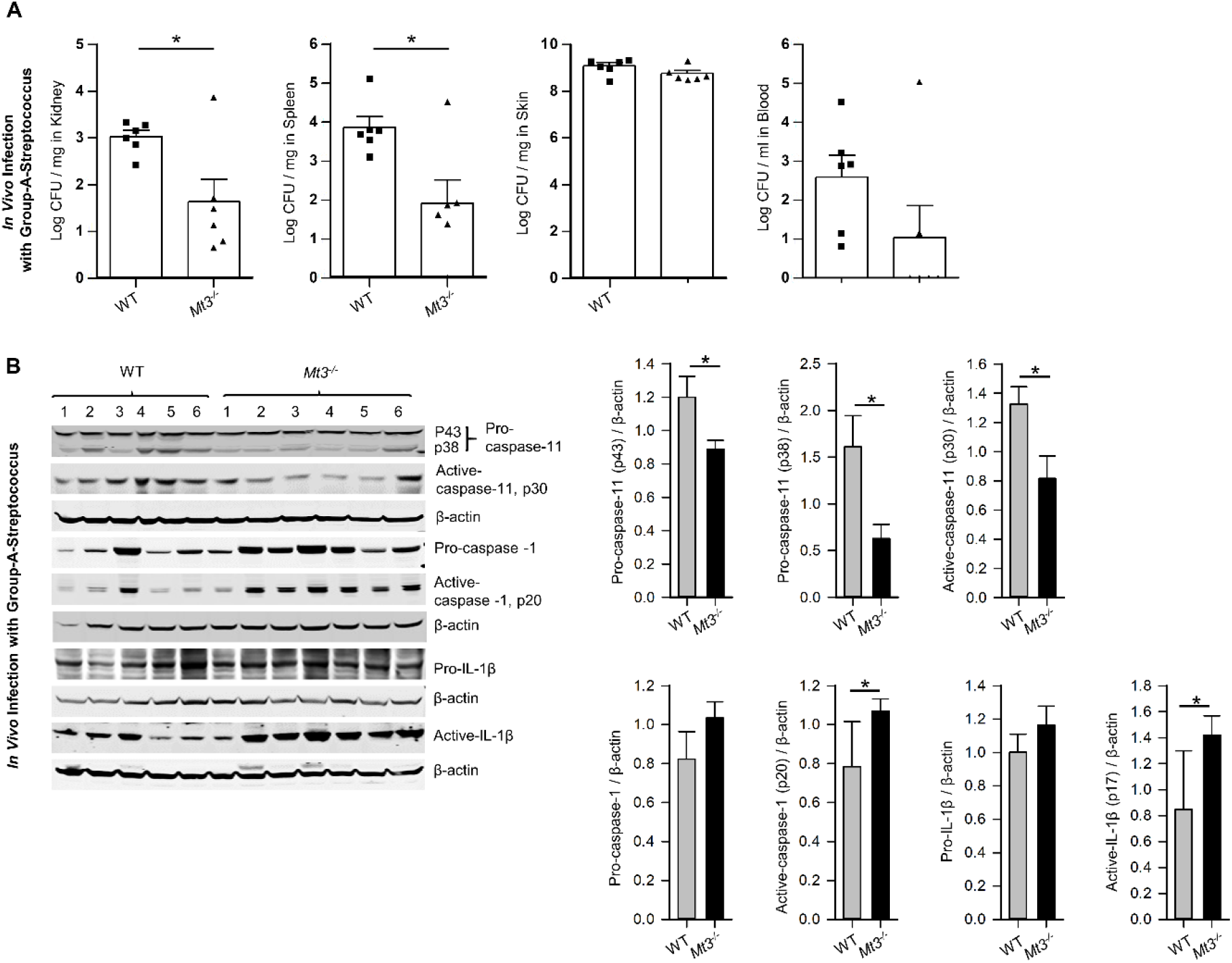
MT3 suppresses caspase-1 and IL-1β activation and antibacterial immunity in Group A Streptococcus infected mouse. See also. **Fig. 2 | A**, WT and *Mt3^-/-^* mice infected *s.q.* with Group A Streptococcus (GAS5448) for 72h. Bacterial growth measured in kidney, spleen, skin and blood. CFUs were log transformed. n = 6 per group, two-tailed t-test. **B**, Western blots of pro-caspase-11, active-caspase-11, pro-caspase-1, active-caspase-1, pro-IL1β and active-IL-1β in whole kidney homogenates. Bar graphs represent densitometric analysis of targets normalized to β- actin. n = 6 per group, two-tailed t-test, data are mean ± SEM.

**Extended Data Fig. 3.**
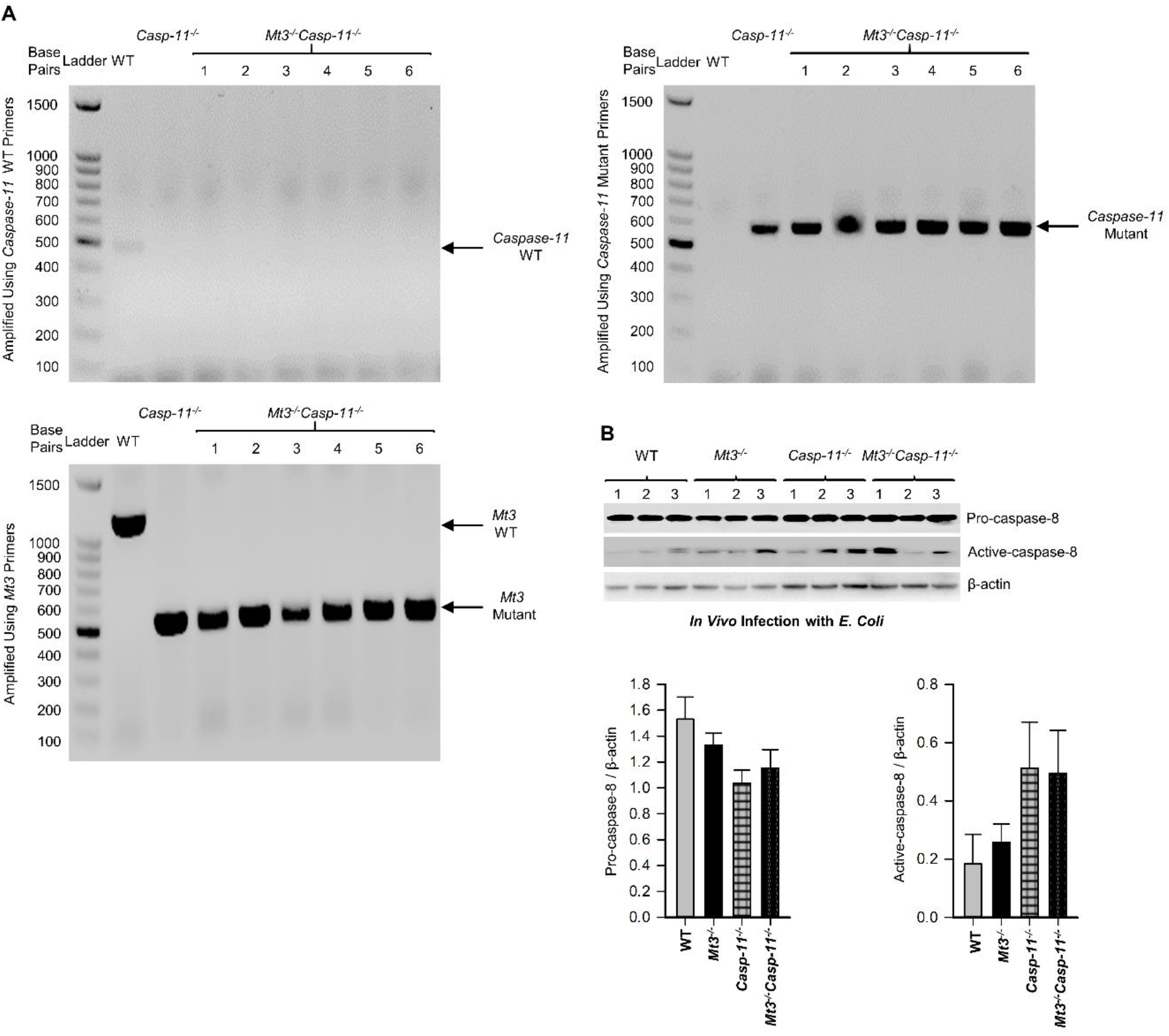
Generation of *Casp11^-/-^Mt3^-/-^* mice and analysis of caspase-8 during *E. coli* infection *in vivo*. See also. **Fig. 3 | A**, *Mt3^-/-^* mice crossed to *Casp4^tm1Yuan^*/J (*Casp11^-/-^*) mice to obtain mice genetically deficient in *Mt3* and *Casp-11* (*Mt3^-/-^Casp11^-/-^*). Tail genomic DNA was amplified to confirm the deletion of *Casp-11* and *Mt3* genes by gel electrophoresis. **B**, Western blots of pro-caspase-8 and active-caspase-8 in kidney homogenates of WT, C*asp-11^-/-^*, *Mt3^-/-^* and *Casp-11^-/-^Mt3^-/-^* mice infected *i.p.* with *E. coli* (1 X10^9^ CFUs/mouse) for 6h. Bar graphs represent densitometric analysis of targets normalized to β-actin, n = 3-6 per group, one-way ANOVA, data are mean ± SEM.

**Extended Data Fig. 4.**
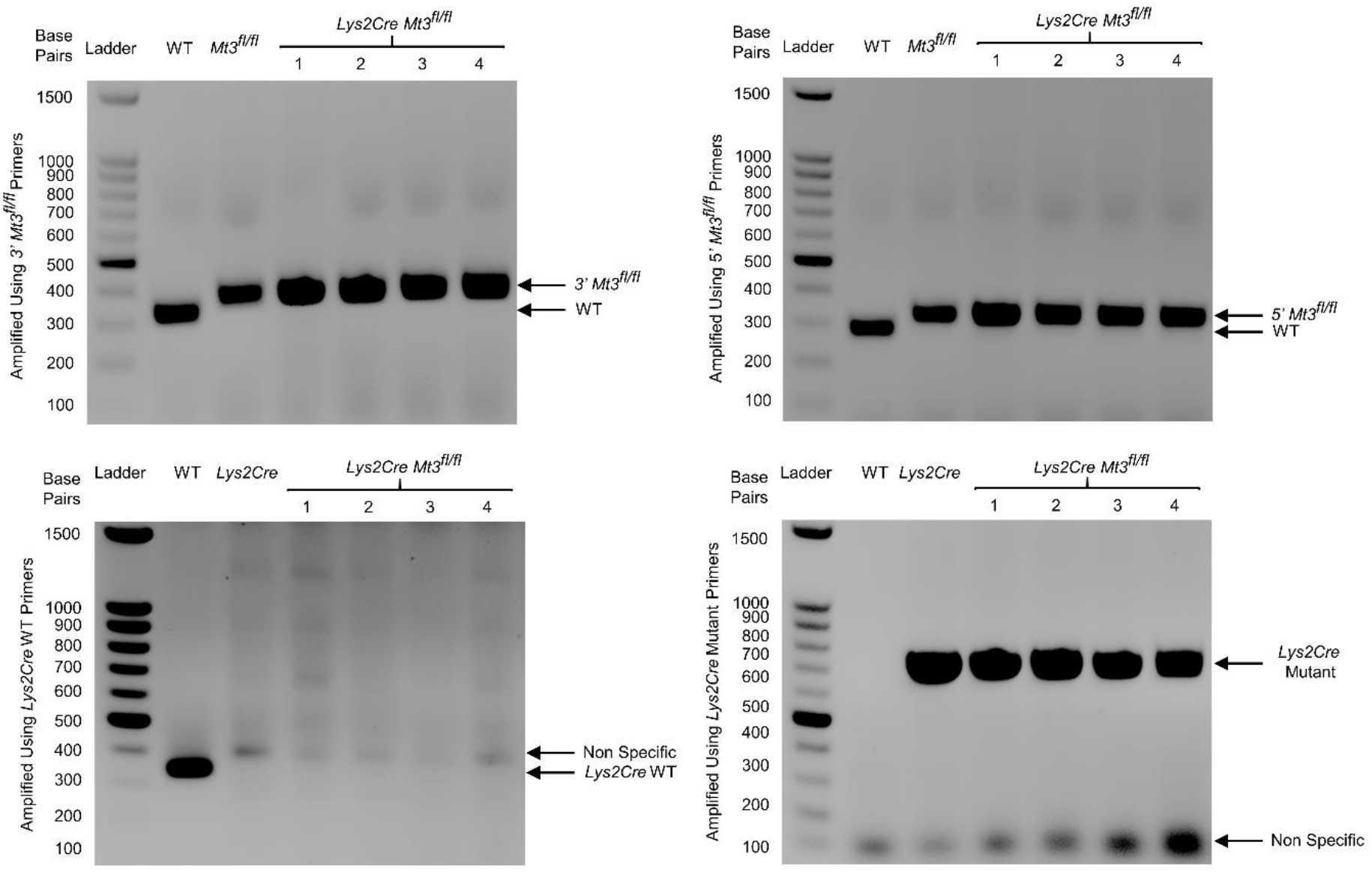
Generation of myeloid-MT3 deficient mice. See also. **Fig. 4 |** Genotyping of WT, *Lys2Cre*, *Mt3^fl/fl^* and *Lys2Cre Mt3^fl/fl^* mice. To confirm 5’ loxp, 3’ loxp and *Lys2Cre* insertion, tail genomic DNA was amplified and analyzed by gel electrophoresis using the primers detailed in the experimental procedures.

**Extended Data Fig. 5.**
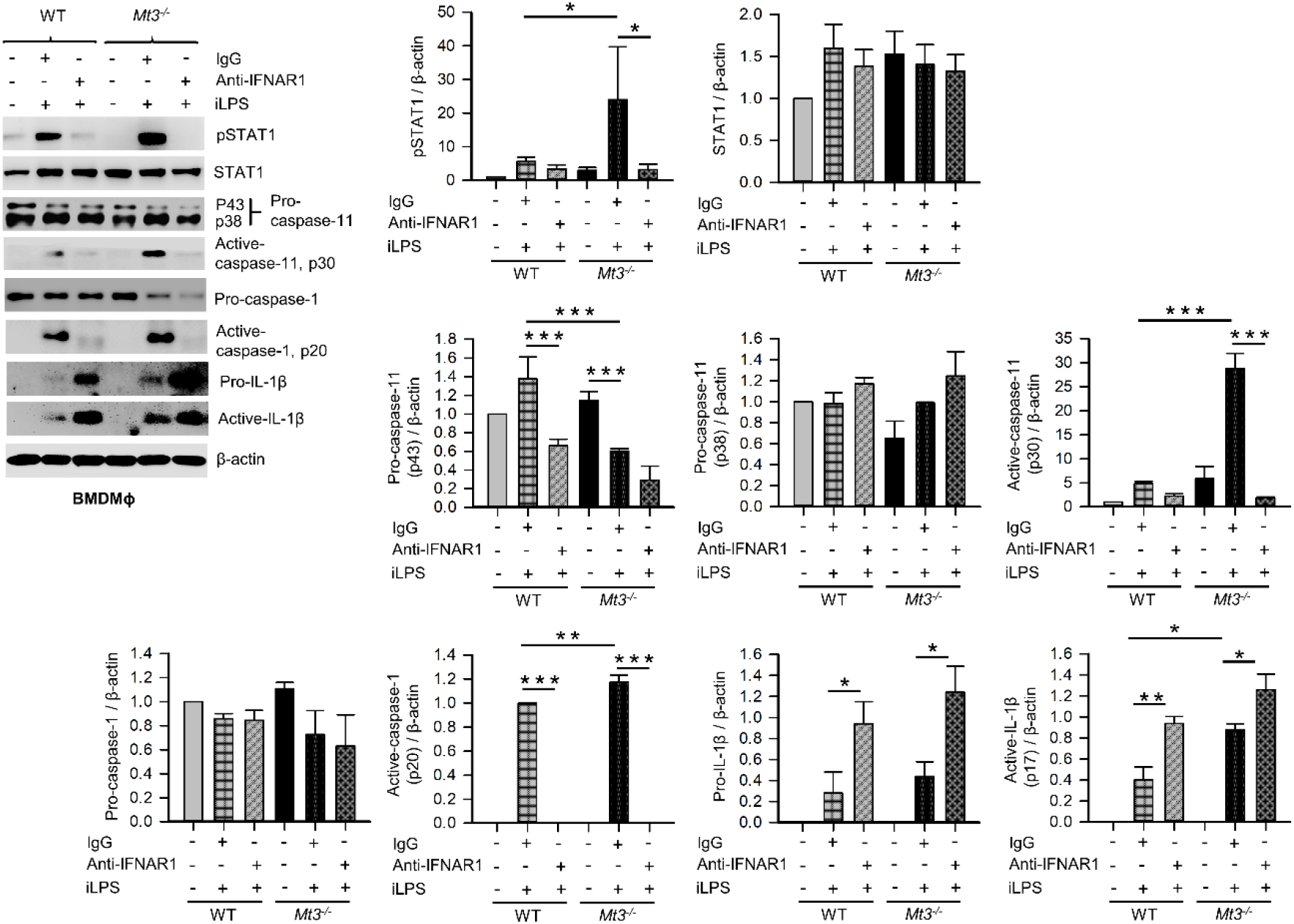
IFNAR1 blockade attenuates caspase-11 and caspase-1 activation, but augments pro- and active-IL-1β levels in BMDMϕ. See also. **Fig. 5 |** WT and *Mt3^-/-^* BMDMϕ treated with monoclonal anti-IFNAR1 or IgG antibody 1h prior to and 24h after iLPS (10 μg/ml, 48h) stimulation. Western blots of pSTAT1, STAT1, pro- caspase-11, active-caspase-11, procaspase-1 and pro-IL1β in cellular extracts and active caspase-1 and active-IL-1β in the cell-free media supernatants. Bar graphs represent densitometric analysis of targets normalized to β-actin. 3 independent experiments, one- way ANOVA, data are mean ± SEM.

**Extended Data Fig. 6.**
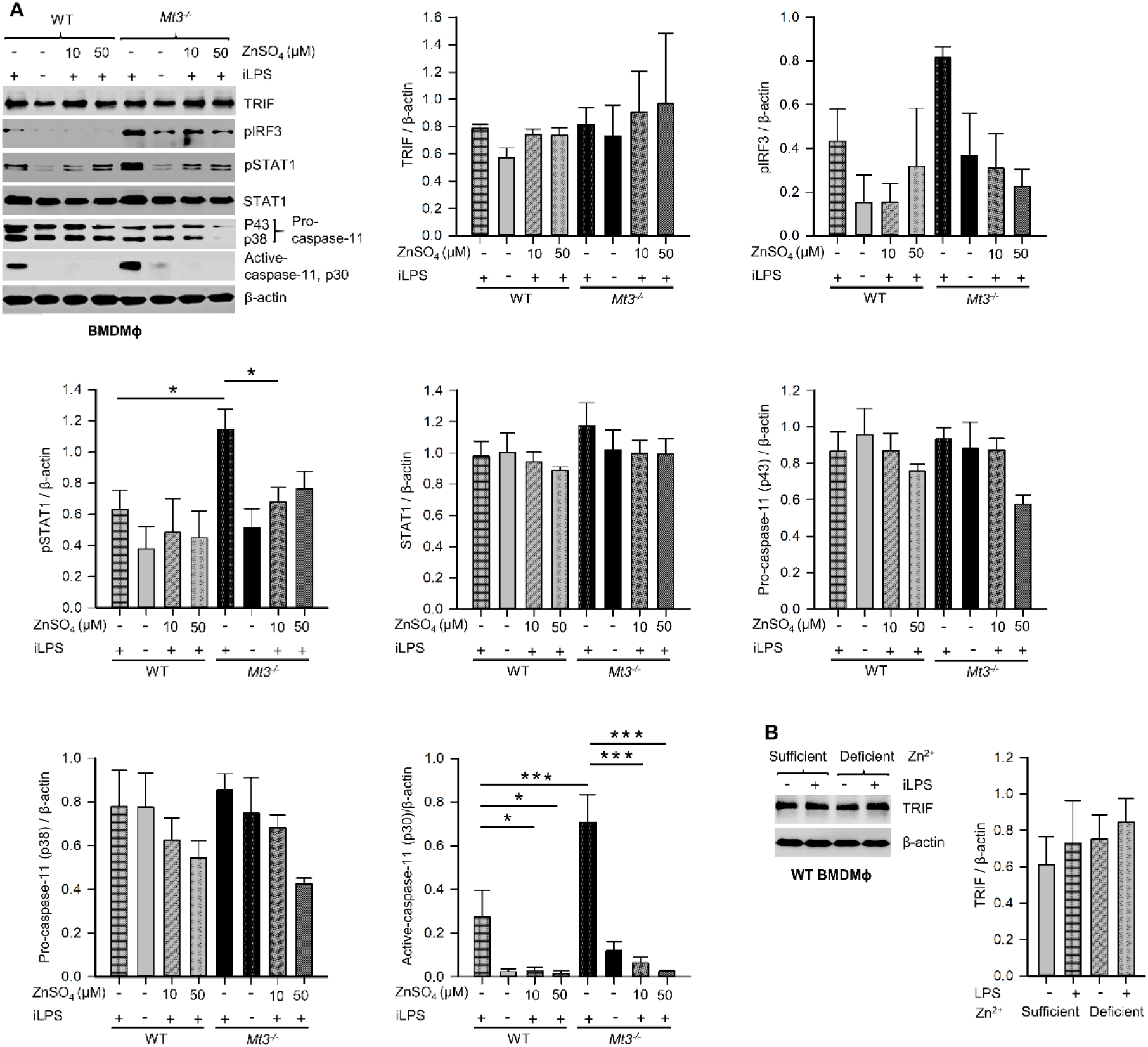
Zn^2+^ supplementation suppresses non-canonical inflammasome activation in WT BMDMϕ. See also. **Fig. 6 | A**, WT and *Mt3^-/-^* BMDMϕ exposed to the indicated doses of ZnSO4 for 3h, followed by stimulation with iLPS (10 μg/ml) or vehicle for 48h. Immunoblots of TRIF, pIRF3, pSTAT1, STAT1, pro-caspase-11 and active-caspase-11 in cellular extracts. Bar graphs represent densitometric analysis of targets normalized to β-actin. 3 independent experiments, one-way ANOVA. **B**, WT BMDMϕ stimulated with iLPS (10 μg/ml) or vehicle for 24h in Zn^2+^- sufficient or Zn^2+^-deficient Opti-MEM media. Immunoblot of TRIF in cellular extracts. Bar graphs represent densitometric analysis of TRIF normalized to β-actin. 3 independent experiments, one-way ANOVA, data are mean ± SEM.

**Extended Data Fig. 7.**
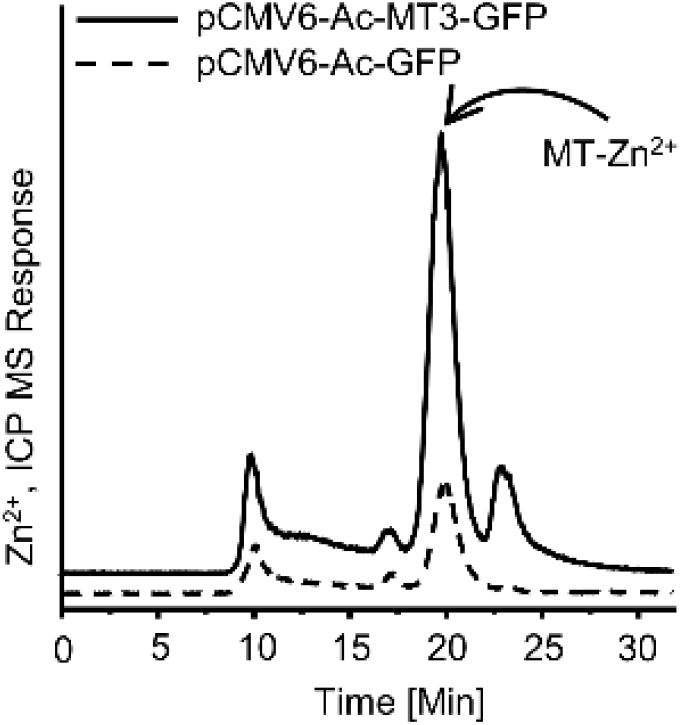
Overexpression of MT3 in *Mt1^-/-^Mt2^-/-^* BMDMϕ. See also. **Fig. 6 |** *Mt1^-/-^Mt2^-/-^* BMDMϕ transfected with the *Mt3* overexpressing vector (pCMV6-Ac-MT3- GFP) or empty vector (pCMV6-Ac-GFP) control for 48 h. SEC-ICP-MS analysis of cell lysates demonstrating an increase in the MT-associated Zn^2+^ signal in cells that were transfected with the pCMV6-Ac-MT3-GFP.

## Methods

### Reagents and resources

| Name | Source | Identifier |
| --- | --- | --- |
| **Antibodies** |  |  |
| anti-TRIF | Proteintech | Cat#23288-1-AP |
| anti-pIRF3 (Ser396) | BIOSS | Cat#bs-3195R |
| anti-STAT1 | Abcam | Cat#ab99415 |
| anti-pSTAT1 (pY701) [M135] | Abcam | Cat#ab29045 |
| anti-GBP5 | Proteintech | Cat#13220-1-AP |
| anti-GBP2 | Proteintech | Cat#11854-1-AP |
| anti-Caspase 11 (17D9) | eBioscience™ | Cat#14-9935-82 |
| anti-Caspase-11 | Abcam | Cat# ab180673 |
| anti-CASPASE-4 | MBL | Cat#M029-3 |
| anti-GSDMD | Proteintech | Cat#20770-1-AP |
| anti-Caspase-1 | AdipoGen Life Sciences | Cat#AG-20B-0042-C100 |
| anti-Caspase-1 (14F468) | Santa Cruz<br>Biotechnology | Cat#sc-56036 |
| anti-IL-1β /IL-1F2 | R&D Systems | Cat#AF-401-NA |
| anti-IL-1β /IL-1F2 | R&D Systems | Cat#MAB4011 |
| anti-IL-1β (B122) | Santa Cruz<br>Biotechnology | Cat#sc-12742 |
| anti-Caspase-8 (1G12) | Enzo Life<br>Sciences | Cat#ALX-804-447-C100 |
| anti-β-actin | Cell Signaling<br>Technology | Cat#4967s |
| anti-β-actin | ThermoFisher<br>Scientific | Cat#PA1-183 |
| anti-β-actin | R&D Systems | Cat#MAB-8929 |
| Mouse anti-armenian hamster IgG-HRP | Santa Cruz<br>Biotechnology | Cat#sc-2789 |
| Goat anti-rabbit IgG(H+L), HRP conjugate | Proteintech | Cat#SA00001-2 |
| IRDye® 800CW goat anti-rabbit IgG | LI-COR<br>Biosciences | Cat#926-32211 |
| IRDye® 680RD goat anti-rabbit IgG | LI-COR<br>Biosciences | Cat#926-68071 |
| Goat anti-mouse IgG(H+L), HRP conjugate | Proteintech | Cat#SA00001-1 |
| IRDye® 680RD goat anti-mouse IgG | LI-COR<br>Biosciences | Cat#926-68070 |
| Mouse IgG (H&L) secondary antibody<br>peroxidase conjugated pre-adsorbed | Rockland<br>Immunochemicals | Cat#610-1319-0500 |
| Rabbit anti-goat IgG(H+L), HRP conjugate | Proteintech | Cat#SA00001-4 |
| Goat anti-Rat IgG(H+L), HRP conjugate | Proteintech | Cat#SA00001-15 |
| IRDye® 800CW goat anti-rat IgG | LI-COR<br>Biosciences | Cat#926-32219 |
| anti-IFNAR1 (Clone: MAR1-5A3) | BioLegend | Cat#127302 |
| anti-IgG1 (Clone: MOPC-21) | BioLegend | Cat#400102 |
| **Bacterial Strains** |  |  |
| *Escherichia coli* (K12) | Dr. Jason Gardner<br>(gardnejr@ucmail.<br>uc.edu) | N/A |
| *Klebsiella pneumoniae* (KP2 2-70) | Dr. Jason Gardner<br>(gardnejr@ucmail.<br>uc.edu) | N/A |
| Group A Streptococcus (*Streptococcus pyogenes*) | Dr. Suba Nookala<br>(suba.nookala@un<br>d.edu) | N/A |
| **Primers** |  |  |
| *Mt3* | Applied<br>Biosystems | Mm00496661_g1 |
| *Mt2* | Applied<br>Biosystems | Mm00809556_s1 |
| *Mt1* | Applied<br>Biosystems | Mm00496660_g1 |
| *Hprt* | Applied<br>Biosystems | Mm00446968_m<br>1 |
| *MT3* | Applied<br>Biosystems | Hs00359394_g1 |
| *HPRT1* | Applied<br>Biosystems | Hs99999909_m1 |
| Genotyping Primers | IDT | N/A |
| **siRNA and expression vector** |  |  |
| *MT3* | Ambion | Cat#AM16708 |
| *Ticam1* | Dharmacon™ | Cat#L-055987-00-0005 |
| ON-TARGETplus™ Control Pool (Non-Targeting pool) | Dharmacon™ | Cat#D-001810-10-20 |
| pCMV6-AC-GFP (PS100010) | ORIGENE | Cat#MG200059 |
| pCMV6-AC-GFP | ORIGENE | Cat# PS100010 |
| **Chemicals and accessories** |  |  |
| RPMI 1640 | CORNING | REF#10-041-CV |
| Opti-MEM® I | GIBCO | REF#31985-070 |
| FBS | HyClone | Cat#SH30396.03 |
| DPBS | CORNING | REF#21-031-CV |
| PBS without calcium and magnesium | CORNING | REF#21-040-CV |
| HBSS | CORNING | REF#21-021-CM |
| HEPES | Sigma | Cat#H3375 |
| Mouse M-CSF | PEPROTECH | Cat#315-02 |
| Human M-CSF | PEPROTECH | Cat#300-25 |
| Trypsin-EDTA | CORNING | REF#25-053-CI |
| LPS-B5 Ultrapure | InVivoGen | Cat#tlrl-pb5lps |
| Glycerol | Fisher Bioreagents | Cat#BP229-1 |
| LB Broth | Fisher Bioreagents | Cat#BP1427-500 |
| BBL™ Brain Heart Infusion Broth | Becton Dickinson | Cat#22182 |
| Agar | BD Bacto™ | Cat#214010 |
| BD Bacto™ Dehydrated Culture Media: Todd<br>Hewitt Broth | BD | Cat#249240 |
| Gibco™ Bacto™ Yeast Extract | Gibco | Cat#212750 |
| BD BBL™ RODAC™ Trypticase™ Soy Agar<br>with 5% Sheep Blood (TSA II) | BD | Cat#221261 |
| EZ Pack™ Agarose | ASi | Item<br>No.#AG2501 |
| ZnSO<sub>4</sub> | Sigma | Cat#z-4750 |
| Sodium Chloride | Sigma | Cat#S1679-1KG |
| Ethyl Alcohol | Fisher Scientific | Cat#A407-4 |
| Methanol | Fisher Scientific | Cat#A434-20 |
| Methanol | BDH | Cat#BDH1135-4LP |
| Dimethyl Sulfoxide | Sigma | Cat#D8418-1L |
| Chloroform | Fisher Scientific | Cat#C5312 |
| Hydrochloric Acid | Fisher Scientific | Cat#A508-212 |
| Sulfuric Acid | Fisher Scientific | Cat#A300<sup>SL</sup>-212 |
| Sodium Hydroxide | Fisher Scientific | Cat#S318-3 |
| Precise Protein Gels | Invitrogen | REF#XP04205B<br>OX |
| SurePAGE™, Bis-Tris, 10x8 gels (4-20%, 15 wells) | GenScript | Cat#M00657 |
| Tris-MOPS-SDS Running Buffer Powder | GenScript | Cat#M00138 |
| Nitrocellulose membranes | BIO-RAD | Cat#162-0112 |
| Protease &Phosphatase Inhibitor Cocktail | Thermo Scientific | Cat#78442 |
| Denaturing Cell Extraction Buffer | Invitrogen | Cat#FNN0091 |
| XCell II™ Blot Module | ThermoFisher<br>Scientifics | Cat#EI9051 |
| Glycine | Fisher Scientific | Cat#BP381 |
| Tris-base | Fisher Scientific | Cat#BP152 |
| SDS | Fisher Scientific | Cat#BP166 |
| Tween 20 | Acros Organics | Cat#23336-0010 |
| Tween 20 | Sigma | Cat#P-1379 |
| Triton X100 | Fisher Scientific | Cat#BP151 |
| Bovine serum albumin | Sigma | Cat#A7030 |
| Intercept® (TBS) Blocking Buffer | LI-COR<br>Biosciences | Cat#927-60001 |
| Non-fat dry milk | Nash Finch Co. | N/A |
| Isoflurane | Covetrus | NDC<br>Code(s)#11695-6777-2 |
| Nair | Chuech & Dwight<br>Co.inc | N/A |
| Gauze | Medline | REF#PRM25444 |
| Puritan cotton tipped applicator | Puritan | SKU#836-WC |
| Insulin injection syringe (1mL) | Exelint<br>international co. | Cat#26029 |
| 10 ml plastic Syringe | Fisherbrand | Cat#14955459 |
| 1 ml plastic Syringe | Fisherbrand | Cat#14955456 |
| BioLite12 well multidish | Thermo Scientific | Lot#<br>H4XA4RE106 |
| 24 well tissue culture plates | CellPro | Lot#<br>072219BA03 |
| 48 well cell culture plate | NEST | Lot#<br>121717A004 |
| Tissue culture plate 96 well, flat bottom | Fisherbrand | Cat#FB012931 |
| 96 well ELISA plate | NEST | Lot#04291818A0<br>07 |
| Petri dish | NEST | Lot#753001 |
| 15 ml Centrifuge tube | Fisher Scientific | Cat#14-955-237 |
| 50 ml Centrifuge tube | Fisher Scientific | Cat#14-955-239 |
| Cell scraper | SPL Life Sciences | Cat#90030 |
| Golden Rod animal Lancet (4mm) | Medipoint Inc | N/A |
| Omni homogenizer Th-01 and tips | Omni International | Model<br>Number#THP115 |
| GeneArt Platinum Cas9 Nuclease | ThermoFisher | Cat# B25641 |
| Alt-R SpCas9 Nuclease 3NLS | ThermoFisher | Cat# 1074181 |
| sgRNA | CCHMC<br>transgenic core | N/A |
| TSK gel 3000SW gel filtration column | TOSOH<br>BIOSCIENCE | Cat#05789 |
| DreamTaq DNA Polymerase | Thermo Scientific | REF#EP0702 |
| **Commercial Assays** |  |  |
| TransIT®-TKO | Mirus Bio LLC | Prod. No.#MIR<br>2150 |
| TransIT®-LT1 | Mirus Bio LLC | Prod. No.#MIR<br>2300 |
| Pierce Protein Transfection Reagent Kit | Thermo Scientific | REF#89850 |
| Human IL-1β/IL-1F2 DuoSet ELISA | R&D Systems | Cat#DY201-05 |
| Mouse IL-1β | BioLegend | Cat#432601 |
| RNeasy Mini Kit | Qiagen | Cat#74104 |
| QUICK-RNA™ MINIPREP KIT | Denville Scientific | Cat#R1055 |
| Reverse Transcription System | Promega | REF#A3500 |
| Probe Lo-Rox 2X qPCR Mix | RADIANT™ | Cat#QP9020 |
| CytoTox 96® NonRadioactive Cytotoxicity Assay kit Promega | Promega | REF#G1781 |
| SuperSignal™ West Femto Maximum Sensitivity Substrate | Thermo Scientific | Cat#34096 |
| NEBNext Poly(A) mRNA Magnetic Isolation Module | New England BioLabs | Cat#E7490L |
| NEBNext Ultra Directional RNA Library Prep Kit | New England BioLabs | Cat# E7420L |
| NEBNext Library Quant Kit | New England BioLabs | Cat# E7630L |
| MEGAshorscript T7 Kit | Thermo Fisher | Cat#AM1354 |
| MEGAclear Kit | Thermo Fisher | Cat# AM1908 |
| Quick-DNA™ Miniprep Plus Kit | ZYMO RESEARCH | Cat#D4069 |
| Radiant™ Taq DNA Polymerase | RADIANT™ | Cat#C101 |

| Experimental Models: Organisms/Strains |  |  |
| --- | --- | --- |
| C57BL/6 Mice | Jackson<br>Laboratory | Stock#00064 |
| C57BL/6 *Mt3*<sup>-/-</sup> deletion of Exon 3 | Transgenic Animal<br>and Genome<br>editing core facility<br>at CCHMC | N/A |
| C57BL/6 *Mt1*<sup>-/-</sup>*Mt2*<sup>-/-</sup> | Dr. George S.<br>Deepe<br>(deepegs@ucmail.<br>uc.edu) | N/A |
| C57BL/6 *Casp4*<sup>tm1Yuan/J</sup> (*Casp-11*<sup>-/-</sup>) | Jackson<br>Laboratory | Stock#024698 |
| *Mt3*<sup>-/-</sup>*Casp-11*<sup>-/-</sup> | Crossed and bred<br>in-house | N/A |
| C57BL/6 *Lys2Cre* | Dr. George S.<br>Deepe<br>(deepegs@ucmail.<br>uc.edu) | N/A |
| C57BL/6 *Lys2Cre Mt3<sup>fl/fl</sup>* | Crossed and bred<br>in-house | N/A |
| **Instrument, Software and Algorithms** |  |  |
| FluorChem® HD2 | Cell Biosciences | S/N#FC HD2<br>Imager |
| Odyssey CLx Imaging system | LI-COR | N/A |
| 7500 Fast Real Time PCR System | Applied<br>Biosystems | S/N#275013253 |
| RNA-seq data analysis - DAVID Bioinformatics<br>Resources v.6.8 | NIAID/NIH | (https://david.ncifcrf.gov/summary.jsp ) |
| Protein-protein interaction networks- STRING<br>v.11.0 | ELIXIR | (https://string-db.org/ ) |
| NIH ImageJ Fiji | NIH | (https://imagej.nih.gov/ij/) |

### Microbes

*E. coli* (K12) and *K. pneumoniae* were kindly provided by Dr. Jason Gardner at the University of Cincinnati. Group A Streptococcus (GAS 5448) was kindly provided by Dr. Suba Nookala at the University of North Dakota.

### Mice

All mice used in this study were on the C57BL/6 background. WT and *Casp4^tm1Yuan^*/J (C*asp-11^-/-^*) mice were acquired from the Jackson Laboratory. *Lys2Cre* mice were kindly provided by Dr. George S. Deepe. Jr. (University of Cincinnati). *Mt3^-/-^* (exon 3 deleted) and *Mt3^fl/fl^* mice were generated using clustered regularly interspaced short palindromic repeats (CRISPR) by the Transgenic Animal and Genome Editing Core facility at the Cincinnati Children’s Hospital Medical Center (CCHMC). In mice, the *Mt* gene cluster is located on chromosome 8. The *Mt3* gene consists of 3 exons and is preceded by *Mt1* and *Mt2* genes. We targeted exon 3 of *Mt3* by flanking it with loxp sites (*Mt3^fl/fl^*) using the CRISPR-Cas9 gene targeting approach. *Lys2Cre Mt3^fl/fl^* mice that exhibit myeloid *Mt3* deficiency were generated by crossing *Lys2Cre* mice to *Mt3^fl/fl^* mice. Deletion of the *Mt3* gene was confirmed in BMDMϕ and PMϕ of *Lys2Cre Mt3^fl/fl^* mice by genotyping as detailed in the CRISPR/Cas9 generation of *Mt3^fl/fl^* mice section. To confirm 3’ loxp site insertion or deletion, genomic DNA amplified using forward (5՛ TAG GCT TCC CAC CTG TTT GG 3՛) and reverse (5՛ GCC AAG ATA AAG TCC GGG GT 3՛) primers. To confirm 5’ loxp site insertion or deletion, genomic DNA amplified using forward (5՛ TCG AAC TAC CTC CAA ACA GAG AAC 3՛) and reverse (5՛ TCA GTT TGG TCC AAA CGG

GAT G 3՛) primers. To confirm *Lys2Cre* gene insertion, genomic DNA amplified using mutant (5՛ CCC AGA AAT GCC AGA TTA CG 3՛), common (5՛ CTT GGG CTG CCA GAA TTT CTC 3՛) and WT (5՛ TTA CAG TCG GCC AGG CTG AC 3՛) primers. *Casp-11^-/-^Mt3^-/-^* mice were generated by crossing *Casp-11^-/-^* mice to *Mt3^-/-^* mice. *Casp-11* was amplified using tail genomic DNA by mutant reverse (5՛ CGC TTC CTC GTG CTT TAC GGT AT 3՛), common forward (5՛ ACA ATT GCC ACT GTC CAG GT 3՛) and WT reverse (5՛ CAT TGC TGA CCT TAT TTC TGT ATG G 3՛) primers. *Mt3* was amplified using tail genomic DNA by forward (5՛ TTG GGG TGA GGT GTA GAG GT 3՛) and reverse (5 GCC AAG ATA AAG TCC GGG GT ՛ 3՛) primers. Mice used in this study had ad-libitum access to food and water. All mice were housed in the Department of Laboratory Animal Medicine, University of Cincinnati, accredited by American Association for Accreditation of Laboratory Animal Care (Frederick, MD) and experiments were conducted in accordance with Animal Welfare Act guidelines of the National Institutes of Health.

### CRISPR/Cas9 generation of *Mt3^fl/fl^* mice

The methods for the design of sgRNAs, donor oligos and the production of *Mt3^fl/fl^* (loxp sites surrounding exon 3 of the murine *Mt3* gene) animals were as described previously ^74^. The sgRNAs were selected according to the on- and off-target scores from the CRISPR design web tool (http://genome-engineering.org) as well as CRISPOR (http://crispor.tefor.net) ^75^. The selected sgRNA target sequences were cloned, according to the published method ^76^, into the pX458 vector (addgene #48138) that was modified by us to contain an optimized sgRNA scaffold ^77^ and a Cas9- 2A-GFP. Their editing activity were validated by the T7E1 assay in mouse mK4 cells ^78^, compared side-by-side with Tet2 sgRNA that was known to work in mouse embryos efficiently ^79^. Validated sgRNA was transcribed *in vitro* using the MEGAshorscript T7 kit (ThermoFisher) and purified by the MEGAclear Kit (ThermoFisher), and stored at -80°C. To prepare the injection mix, we incubated sgRNA and Cas9 protein (ThermoFisher) at 37°C for 5 mins. to form the ribonucleoprotein complex and then added the donor oligos to it. The initial attempt was to insert both loxP sites simultaneously via piezo-driven cytoplasmic injection ^80^ of 100 ng/ul Cas9 protein, 50 ng/ul 5’ sgRNA, 50 ng/ul 3’ sgRNA, 50 ng/ul 5’ donor, and 50 ng/ul 3’ donor into fertilized eggs. Injected eggs were transferred into the oviductal ampulla of pseudo-pregnant CD-1 females on the same day. Pups were born and genotyped by PCR and Sanger sequencing. However, only 3’ loxP-containing mice were obtained from this attempt. After breeding them to homozygosity for the 3’ loxP, a new set of 5’ sgRNA and the donor oligo was designed and injected into the zygotes with the mix containing 150 ng/ul Cas9 protein, 75 ng/ul 5’ sgRNA, and 100 ng/ul ssDNA donor oligo. Injected eggs were transferred into the oviductal ampulla of pseudopregnant CD-1 females on the same day. Pups were born and genotyped by PCR and Sanger sequencing. Founder mice carrying both 5’ and 3’ loxP sites in cis were finally obtained. Animals were housed in a controlled environment with a 12-h light/12-h dark cycle, with free access to water and a standard chow diet. All animal procedures were carried out in accordance with the Institutional Animal Care and Use Committee-approved protocol of Cincinnati Children’s Hospital and Medical Center.

### Mϕ culture

hMϕ were prepared from peripheral blood mononuclear cells (PBMCs). Briefly, human blood obtained from the Hoxworth Blood Center, University of Cincinnati was diluted (1:2) with calcium- and magnesium-free 1X Dulbecco’s phosphate-buffered saline (DPBS) and inverted gently to mix. Ficoll-Paque (10 ml for the total volume of 40 ml diluted blood) was layered at the bottom of the tube. The tubes were centrifuged at 400 X g for 30 mins. without break at 20°C. The PBMC interface was transferred to sterile tubes and washed three times using 40 ml of DPBS containing 2mM EDTA and centrifuged at 120 X g for 10 mins. without break at 4°C. A final wash was performed with DPBS (without EDTA). Isolated PBMCs were resuspended in complete RPMI 1640 medium. PBMCs (5 × 10^6^) were plated in 24 well plates containing complete RPMI medium. After 24h, adherent monocytes were washed three times with DPBS and plated in complete RPMI 1640 medium (Corning®) containing 10 ng/ml macrophage-colony stimulating factor (M-CSF), 10%FBS, 10 μg/ml gentamycin sulfate (Alkali Scientific Inc.) and 2-mercaptoethanol. Cells were differentiated by exposure to human recombinant M-CSF on days 0, 2 and 4. After 6 days, hMϕ were washed with DPBS prior to use for the experiment.

Mouse BMDMϕ were prepared by differentiating bone marrow cells in complete RPMI 1640 medium containing 10 ng/ml mouse M-CSF, 10% fetal bovine serum (FBS) (HyClone Laboratories, Utah), gentamycin sulfate (10 μg/ml) and 2-mercaptoethanol. BMDMϕ were fed on days 0, 2 with complete RPMI 1640 medium containing 10 ng/ml M-CSF and were supplemented on day 4 with M-CSF. After 6 days, adherent Mϕ were harvested by washing with DPBS followed by trypsinization and centrifuged at 1600 rpm for 5 mins. at 4°C. Mϕ were washed again with DPBS at 1600 rpm for 5 mins. at 4°C and counted under the microscope. Dead cells were excluded from enumeration using Trypan Blue stain. BMDMϕ (0.5 × 10^6^ in 24 well or 1 × 10^6^ in 12 well plate) were seeded.

### Bioinformatics analysis

The STRING database was used to review the protein-protein interaction networks of MT3 in *Mus musculus* and *Homo sapiens* ^22^. A full network analysis was conducted based on text mining, experiments, databases, co-expression, neighborhood, gene fusion and co-occurrence data with a minimum required interaction score of 0.4 and a maximum of 50 interactors in the first shell and 50 interactors in the second shell. Statistical significance of the enriched biological processes (GO BP categories) in the MT3 network was set with a false discovery rate (FDR) <0.05.

Identification of differentially expressed genes in resting WT compared to resting *Mt3^-/-^* Mϕ was based on our previously published RNA-seq data (NCBI SRA: PRJNA533616) ^19^. The number of biological replicates used in the analysis was 3 per group. Genes differentially expressed with a fold change FC>2 and adjusted p value q<0.05 were considered significant. The Benjamini-Hochberg correction was used to adjust p values for multiple hypothesis testing. Differentially expressed genes in *Mt3^-/-^* BMDMϕ compared to WT BMDMϕ with q<0.05 were queried using the functional annotation clustering tool DAVID to identify statistically enriched GO categories (using the GO terms, BP direct, CC direct and MF direct) in *Mt3^-/-^* BMDMϕ compared to WT BMDMϕ ^37^. Significance of enrichment was set to FDR <0.05.

### Gene silencing

For gene silencing, Mϕ were transfected with the transfection complex (50 μl) of siRNA and TransIT-TKO® (0.5 %) transfection reagent (Mirus Bio™) in 500 μl of complete RPMI 1640 medium without antibiotics as per the manufacturer’s instructions. Concentration of siRNAs used for gene silencing were 100 nM each of the non-targeting pool (ON-TARGETplus^tm^ scramble siRNA), human *MT3* (MT3 Silencer^®^ Pre-designed siRNA) and mouse *Ticam1* (ON-TARGETplus SMARTpool). All siRNAs were purchased from Dharmacon (GE Healthcare). Both BMDMϕ and hMϕ were incubated with the siRNA containing transfection complexes for 24h in RPMI medium and washed prior to transfection with LPS in Opti-MEM medium. TRIF silencing was assessed by protein expression using Western blots. Human *MT3* silencing was assessed by gene expression using qRT-PCR.

### IFNAR1 neutralization

IFNAR1 on WT and *Mt3^-/-^* BMDMϕ was neutralized using 10 μg/ml monoclonal anti-IFNAR1 antibody (BioLegend; Clone: MAR1-5A3) 1h prior and 24h after iLPS (10 μg/ml) stimulation. Negative control groups were treated with the same dose of isotype control IgG antibody (BioLegend; Clone: MOPC-21). After a total 48h, cell lysates and supernatants were harvested for the molecular analysis.

### Non-canonical inflammasome activation in Mϕ

To activate the non-canonical inflammasome, 1 × 10^6^ Mϕ were transfected with a transfection complex (50 μl) of 0.3% TransIT™-LT1 (Mirus Bio™) transfection reagent (vehicle) and 2 μg/ml or 10 μg/ml ultrapure LPS-B5 (InvivoGen) prepared from *E. coli* 055:K59(B5) in 500 μl Opti-MEM medium (Thermo Fisher Scientific-US) as per manufacturer’s instructions for 24h or 48h.

### Preparation of Zn^2+^-sufficient and Zn^2+^-deficient Opti-MEM media

Molecular biology grade chelex-100 resin (BioRad) was washed three times with metal free ddiH2O prior to use. To prepare Zn^2+^-deficient Opti-MEM medium, washed Chelex-100 resin (3 g per 100 ml) was added to Opti-MEM media and vigorously shaken for 1h on an orbital shaker at room temperature. After this time, media was filtered using a 0.22 μm filter and mixed with fresh washed chelex-100 resin and the same procedure was repeated for a total of 3 times to eliminate metals from Opti-MEM media. Chelex was removed from the media by a final filtration step. During each of these stages, an aliquot of the media was saved to monitor the efficiency of Ca^2+^, Mg^2+^, Mn^2+^, Co^2+^, Zn^2+^, Cu^2+^, Ni^2+^ and Fe^2+^ elimination by ICP-MS. The amount of Zn^2+^ in Opti-MEM media was decreased by 95% by the above chelation method. To prepare Zn^2+^-sufficient media, chelexed Opti-MEM was reconstituted with Ca^2+^, Mg^2+^, Mn^2+^, Co^2+^, Cu^2+^ and Zn^2+^ at the original concentrations as measured by ICP-MS. To prepare Zn^2+^-deficient media, all measured elements except Zn^2+^ were added to the chelexed Opti-MEM media at the original measured concentrations. Finally, the pH of Zn^2+^-sufficient and Zn^2+^-deficient Opti-MEM media was adjusted to 7.4 and filtered prior to use.

### MT3 overexpression and purification

The pCMV6-Ac-GFP vector containing the mouse *Mt3* gene (pCMV6-Ac-MT3-GFP) and empty pCMV6-Ac-GFP vectors were acquired from Origene and dissolved in nuclease-free sterile H2O. Plasmid DNA (5 ng) was added to 50 μl of thawed Novablue competent *E. coli* cells (EMD Millipore) and transformation was performed as per manufacturer’s instructions. *E. coli* cells were serially diluted in S. O. C media (ThermoFisher Scientific) and plated onto Luria-Bertani

(LB) plates with 50 g/ml carbenicillin and grown for 24h at 37°C. A single colony was isolated and inoculated in LB media containing carbenicillin and grown at 37°C for 5h in a shaker. The culture was further amplified by passaging for another 24h. *E. coli* cells were then harvested by centrifugation at 2000 rpm for 10 mins. and plasmid was extracted using the EndoFree plasmid MAXI kit (Qiagen) as per the manufacturer’s instructions. The plasmid was reconstituted in endotoxin-free TE buffer and OD readings obtained were in the range of 1.8-1.9. The resulting endotoxin-free plasmid DNA was set to a concentration of 1 mg/ml in filter-sterilized EndoFree TE buffer and frozen into aliquots until further use.

*Mt1^-/-^Mt2^-/-^* BMDMϕ were transfected with pCMV6-Ac-GFP control vector or pCMV6-Ac-MT3-GFP vector using the LT1 transfection reagent (Mirus Bio) in RPMI media containing 10% serum without antibiotics as per the manufacturer’s instructions. After 48h, BMDMϕ cultures were lysed with 250 μl of 0.1% SDS prepared in double-deionized (ddi) H2O for 20 min. on ice with intermittent mixing. Cell lysates were transferred to 0.22 μm filter tubes and centrifuged at 13000 rpm for 5 mins. Filtered cell lysates were subjected to SEC-ICP-MS to isolate the MT3 protein as described below.

### Preparation of apo-MT3, 4Zn^2+^-MT3 or 6Zn^2+^-MT3

Cell lysates from above were analyzed by SEC-ICP-MS to detect the MT3-associated peak, followed by collection of the fraction of interest (18-21 mins.). The collected fraction was concentrated by freeze drying in Millrock lyophilizer (Millrock, NY). The concentrated fraction was treated with 1 g of Chelex X-100 resin to remove the divalent metals associated to the protein. The total MT concentration was calculated by the total sulfur concentration in the sample with a 1:20 stoichiometry. The sample was divided into 3 fractions, and each fraction was incubated for 2 h in 50 mM Tris-HCl with the appropriate concentration of ^66^Zn^2+^ nitrate to obtain an MT3-Zn^2+^ saturation of 0, 4 or 6 Zn^2+^ ions per MT3 molecule. After incubation, samples were filtered using a 3 kDa MWCO filter to remove the unbound Zn^2+^ and reconstituted in 1X PBS.

### Transfection of apo, Zn^2+^-MT3 complexes into BMDMϕ

In a 24 well plate, 5 × 10^5^ *Mt3^-/-^* BMDMϕ were transfected with the transfection complex (10 μl) containing 500 ng of apo-MT3, 4Zn^2+^-MT3 or 6Zn^2+^-MT3 and Pro-Ject^TM^ (1.75 μl) protein transfection reagent (ThermoScientific-US) in 250 μl of 2% FBS containing antibiotic free RPMI 1640 medium. Control *Mt3^-/-^* BMDMϕ were treated with Pro-Ject^TM^ alone. Cells were incubated with the transfection complexes for 3.5h and washed two times using HBSS prior to challenge with iLPS in Zn^2+^ free Opti-MEM medium. At the experiment end point, cell lysates were either prepared for SEC-ICP-MS analysis or both cell lysates and supernatants were collected for the analysis of non-canonical inflammasome activation.

### SEC-ICP-MS-MS analysis and normalization of data

SEC-ICP-MS-MS analysis of WT and *Mt3^−/−^* BMDMϕ was performed as described previously ^11^. Mϕ were plated in Opti-MEM media, and either left untreated or transfected with 2 μg/ml LPS for 1, 24 and 48h. After this time, Mϕ were washed twice in HBSS and cells were lysed with 0.1% SDS on ice for 20 mins. Cell lysates were then centrifuged in 0.22 mm filter tubes at 13000 rpm for 10 mins. Filtered cell lysates were frozen at -80°C until further analysis by SEC-ICP-MS. 50-80 µl of cell lysates were injected to the HPLC-ICP-MS system according to protein concentration. To normalize the response of ICP-MS-MS signal from SEC separations on different days, 50 μl of 0.5 mg/ml carbonic anhydrase was injected into the liquid chromatography system, and area of Zn^2+^ signals from samples was normalized to area of the carbonic anhydrase peak from each day. The absorbance of carbonic anhydrase at 280 nm was followed to ensure protein integrity.

The instrumentation consisted of an Agilent 1100 HPLC equipped with a degasser, a binary pump, a thermostated auto sampler, a column oven compartment and a diode array detector. For the Mϕ lysates, a TSK gel 3000SW gel filtration column (TSK Tokyo Japan) 7.8 × 300 mm, 10 mm particle size was used. The mobile phase was ammonium acetate pH 7.4, 0.05% MeOH at 0.5 ml/minute. The HPLC system was coupled to the ICP-MS-MS nebulizer via a short polyether ether ketone capillary of 0.17 mm internal diameter. An Agilent 7500ce ICP-MS system equipped with a micromist quartz nebulizer, a chilled double pass Scott spray chamber and a standard 2 mm insert quartz torch with shield torch was used for all experiments. The ICP-MS was operated by the Agilent Mass Hunter integrated chromatographic software in the helium collision mode as reported previously ^11^. The isotope dilution experiments were processed by exporting the chromatographic data to Origin (Origin labs, CA) and the signal, in the form of counts per second, was used to calculate the ratio of ^66^Zn^2+^ / ^64^Zn^2+^ at every point in the chromatograms. This was used to generate a new chromatogram that reflected the input of ^66^Zn^2+^ from the MT3-Zn^2+^ complex at every molecular mass region in the chromatogram. The total area under the chromatograms was used to calculate the concentration of total Zn^2+^ (^64^Zn^2+^) and ^66^Zn^2+^ from the MT3-^66^Zn^2+^ complexes (^66^Zn^2+^ / ^64^Zn^2+^) against a calibration curve of Zn^2+^ based on carbonic anhydrase.

### ICP-MS-MS and SEC-ICP-MS-MS quality control to avoid external Zn^2+^ contamination

All metal analysis experiments were performed using trace metal grade reagents with acid washed plastic vials. Reagent blanks were used to correct the background signal. The analysis was performed through a metal free encased auto sampler. The concentration of Zn^2+^ in the blanks was always below 100 parts per trillion (ppt), the blank estimate concentration on the calibration curves was always below 50 ppt, while the detection limits were below 30 ppt.

For chromatographic analysis, the mobile phase was cleaned using a Chelex 100 resin, using the batch method. In brief, 3 g of Chelex-100 was added to a liter of mobile phase, stirred for 30 mins. and passed through a 0.45 mm membrane. This decreased the Zn^2+^ concentration below 200 ppt (measured as total). By this method, the base line ICP-MS-MS ^66^Zn^2+^ signal was below 1000 counts per second, which represents sub-ppb levels. The SEC column was cleaned using 10 volumes of 0.2 M NaCl and equilibrated with the mobile phase, followed by injection of 50 μl of 2% HNO3 three times to remove any accumulated Zn^2+^ in the column. With this procedure, Zn^2+^ distribution in the samples never deviated more than 10% compared to the theoretical natural Zn^2+^ isotope distribution in the control Mϕ samples. Four blanks and four carbonic anhydrase standards were injected after the cleaning procedure for monitoring Zn^2+^ signal by ICP-MS-MS to ensure optimal column performance. Typically, the column was cleaned every 30-40 samples.

### LPS treatment *in vivo*

Thirteen-week-old WT and *Mt3^-/-^* mice were primed with *i.p.* injection of 10 mg/kg poly(I:C) for 6h followed by 2 mg/kg ultrapure LPS-B5 (InvivoGen) prepared from *E. coli* 055:K59(B5) (*i.p.* injection) for 18h. At the experiment end point, blood was collected by cardiac puncture, allowed to clot, and centrifuged at 2000 rpm for 30 mins. at 4°C to isolate serum. Serum was used to measure cytokines by enzyme-linked immunosorbent assay (ELISA).

### *In vitro* and *in vivo* infection with gram-negative bacteria

For *in vitro* infection, *E. coli* (K12) was grown in LB broth at 37°C overnight in an orbital shaker. The culture was pelleted, washed and resuspended with ice-cold DPBS. Optical Density (OD) of the culture was measured at 600nm using a spectrophotometer. To analyze *E. coli* burden in *in vitro*, hMϕ were transfected with scramble siRNA or *MT3* siRNA as described above and infected with a multiplicity of infection (MOI) of 25 *E. coli*: 1 hMϕ for 3.5h in Opti-MEM medium. hMϕ were washed 3 times with 10 μg/ml gentamycin sulfate containing DPBS to kill extracellular bacteria and incubated in Opti-MEM media with antibiotic for 24h. hMϕ were again washed 3 times with antibiotic-free DPBS, diH2O was added and cells were incubated for 30 mins. to induce osmotic lysis. Cells were scraped and lysates were diluted in DPBS followed by plating on LB agar plates and incubated at 37°C for 24h. Colonies were enumerated as above. Intracellular bacterial burden was represented as percent inhibition of bacterial growth in *MT3*-silenced hMϕ compared to scramble siRNA treated hMϕ.

To analyze antibacterial immunity *in vivo*, 10 to12 week-old mice were used. Mice were infected with *E. coli* 1 × 10^9^ CFUs via *i.p.* injection (300 µl/mouse) for 1h or 6h. *K. pneumoniae* KP2 2-70, a virulent, heavily encapsulated gram-negative bacterial strain ^81, 82^, was grown overnight in brain heart infusion (BHI) broth. The following morning bacteria were washed with DPBS, and administered at 4 × 10^4^ CFUs in 50 µl per mouse by the *i.n.* to isoflurane-anesthetized mice for 48h. At the infection end point, blood was collected by cardiac puncture. A portion of the blood sample was acquired in anticoagulant (3% Na-citrate or EDTA) containing tubes to determine bacterial CFUs in blood. The remaining blood was allowed to clot, and centrifuged at 2000 rpm for 30 mins. at 4°C to isolate serum. Peritoneal lavage was collected using ice-cold 10 ml DPBS. Kidney, lung, and spleen were collected after perfusion with 3 ml of DPBS, indicated organs and skin was rinsed in DPBS and ground with 5 ml DPBS using a glass grinder. Bacterial growth was measured in blood, peritoneal lavage, kidney, lung, skin and spleen samples. Serum and peritoneal lavage were used to measure cytokines by enzyme-linked immunosorbent assay (ELISA).

### *In vivo* infection with gram-positive bacteria

A representative M1T1 clonal Group-A- Streptococcus GAS5448 was used for subcutaneous infections ^83^. GAS was grown at 37°C under static conditions in Todd-Hewitt broth (BD, MD, USA) supplemented with 1.5% yeast extract (BD Biosciences, MD, USA) and *in vivo* infections were performed as described previously ^25, 84^. WT and *Mt3^-/-^* mice (n=8 / group) were used in this study as a model for subcutaneous GAS infections. One day prior to infection, the hair on the back of the mice was depilated (using Nair cream) and mice were infected subcutaneously with

0.1 ml of GAS suspension prepared in sterile DPBS (Ca^2+^/Mg^2+^ free, low endotoxin, Mediatech, VA, USA, DPBS) (OD600 adjusted to yield ∼1-5x10^8^ CFUs). Actual inoculum was determined by plating on trypticase soy agar containing 5% sheep blood (BD Biosciences, MD, USA). Mice were monitored twice daily for body weight, lesions, and mortality. To determine GAS dissemination and load, mice were humanely euthanized 72h post-infection. Blood was drawn through cardiac puncture; necrotic skin, kidney, and spleen were recovered aseptically and weighed. One ml of DPBS was added per 100 mg tissue and homogenized (Omni International, Marietta, GA) followed by plating of ten-fold dilutions on blood agar plates. GAS burden was calculated as colony-forming units (CFUs) per ml (blood) or per mg of tissue. The remaining homogenates were centrifuged for 15 mins. at 12,000 x g at 4°C, and supernatants were stored at -80°C for western blot analysis.

### Gene expression

RNA was isolated from Mϕ after elimination of genomic DNA using RNeasy Plus Mini kit (Qiagen) or QUICK-RNA™ MINIPREP KIT (Thomas Scientific). cDNA was prepared using Reverse Transcription Systems Kit (Promega, WI) or rAmp

First Strand cDNA Synthesis Flex Kit (Thomas Scientific). Taqman primer/probe sets (Applied Biosystems, CA) were used for real-time gene expression analysis using ABI Prism 7500. For time course analysis of expression of murine *Mt* genes, Mϕ were left unstimulated or stimulated with 2 μg/ml iLPS for 0h, 1h, 6h, 24h and 48h. Data are presented as fold change in gene expression normalized to unstimulated Mϕ at the 0h time point. Hypoxanthine guanine phosphoribosyl transferase (*Hprt*) was used as an internal control to compare target gene expression.

### Western blotting

TRIF (Proteintech), pIRF3 (BIOSS), STAT1, pSTAT1 (Abcam), GBP2, GBP5 (Proteintech), caspase-11 (Abcam and eBioscience™), CASPASE-4 (MBL), Gasdermin D (Proteintech and Cell Signaling Technologies), caspase-1 (AdipoGen Life Sciences), IL-1β (R&D Systems) and caspase-8 (Enzo Life Sciences) were assessed in kidney homogenates of *E. coli* infected mice. Cell lysates were prepared using Denaturing Cell Extraction Buffer (Invitrogen) containing protease & phosphatase inhibitor cocktail (ThermoScientific). Culture supernatants were frozen at -80°C until use and processed using methanol-chloroform protein extraction method. Briefly, supernatants were mixed with equal volume of 100% ice-cold methanol and 0.25 times of the total volume of chloroform followed by gentle vortexing and centrifugation at 20,000 X g at 4°C for 10 min. Upper-phase was discarded without disturbing inter-phase proteins. Ice-cold methanol (500 μl) was added to the tube, gently vortexed and centrifuged at 20,000 X g at 4°C for 10 mins. Supernatants were discarded and pellet was dried at 37°C for 3-5 mins. Urea (8 M, pH-8.0) was used to dissolve the pellet and extracted proteins were stored at -80°C. Total cell lysates, supernatants proteins and kidney homogenates were boiled in SDS-PAGE 1X sample buffer at 95°C for 5 mins. Kidney homogenates were centrifuged at 20,000 X g at 4°C for 15 mins. Supernatants were collected for protein analysis. Reduced proteins were run on 8%,10% or 12% SDS-PAGE gels and transferred on to 0.22 μm nitrocellulose membranes (GE Healthcare Life Sciences). Membranes were blocked using 5% skim milk in 1X Tris-buffered saline and 0.1% Tween 20 (1XTBST) and probed overnight with primary antibodies at 4°C. Membranes were washed 3 times for 10 mins. each with 1XTBST and probed with corresponding HRP conjugated or IRDyes (LI-COR) secondary antibodies, washed and developed using BrightStar^TM^ Femto HRP Chemiluminescent 2-Component Substrate Kit (Alkali Scientific Inc. β-actin was used as an internal loading control. Western blots were analyzed using ImageJ software and densitometry data were normalized to β-actin.

### ELISA

Human and mouse IL-1β (BioLegend) concentration in media supernatants, serum and in peritoneal lavage were determined using commercial ELISA kits according to the manufacturer’s instructions.

### Quantification and statistical analysis

Data were analyzed using Sigma plot or GraphPad Prism by one-way ANOVA for multiple comparisons using the indicated ad-hoc methods with at least 3 or more independent biological replicates. Where two groups were compared, two-tailed t-test was used. For *in vivo* infection, bacterial CFUs were log-transformed for statistical analysis. p values were calculated, *p < 0.05, **p < 0.01, ***p< 0.001; NS, not significant, ND, not detected.

